# Inferring structure factors of weakly populated excited states in perturbative crystallography experiments

**DOI:** 10.64898/2026.04.16.719053

**Authors:** Andrew K. Choe, Harrison K. Wang, Doeke R. Hekstra

## Abstract

Perturbative X-ray crystallography can visualize functional dynamics and conformational changes in proteins at atomic resolution. During a typical perturbative crystallography experiment, only a fraction of protein molecules in a crystal will be perturbed, or “excited”. As a result, the observed data represent a mixture of excited and ground states. The conventional approach to estimating the excited-state structure factor amplitudes is to linearly extrapolate the difference between the structure factor amplitudes of the perturbed and unperturbed data. This approach often fails to yield well-refined structural models because it amplifies experimental errors and neglects phase differences between the ground and excited states. Here, we introduce an approach to estimating excited-state structure factor amplitudes that starts from a statistical prior for the correlations between excited and ground states. Using benchmarks from time-resolved crystallography and a drug-fragment screen, we illustrate how this approach effectively addresses the limitations of traditional extrapolation.

## I. INTRODUCTION

Perturbative X-ray crystallography experiments visualize proteins in both their ground and excited states. Comparison of the resulting structures can identify energetically accessible and functionally relevant motions, shedding light on the mechanics of protein function at atomic resolution. For example, in time-resolved X-ray crystallography (TR-X), diffraction is measured on unperturbed crystals, as well as crystals during or after perturbation. Established perturbations for TR-X include light^1–3^, electric field^4–6^, and, in the case of enzyme crystals, their known substrates^7–9^. Other perturbative crystallography methods involve comparing protein constructs that have been crystallized in distinct states or under different conditions. An application of this approach is drug-fragment screening, in which protein crystals are soaked with small molecules from a large library of candidate ligands^10,11^. Solving the resulting structures can identify both the small molecules that bind to the target protein and the resulting structural changes they induce^12,13^.

For both kinds of perturbative crystallography experiments, a fundamental challenge is that the perturbation typically does not excite all protein molecules within the crystal. As such, the perturbed electron density represents a mixture of ground and excited states^14–16^, known as partial excitation. A model of partial excitation in a perturbative crystallography experiment illustrates this concept and its consequent challenges. Consider a theoretical crystal that can be fully excited. Let *V* be the unit cell volume of the crystal. Denote the ground-state electron density as *ρ*_GS_ and the fully excited electron density as *ρ*_ES_. Assume these states are isomorphous. Then, during an experiment where the crystal is only partially excited, the observed electron density *ρ*_ON_ at real-space coordinate **x** = (*x, y, z*) is^17^:

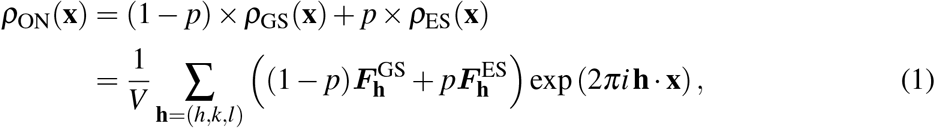

where *p* ∈ [0, 1] is the excited-state fraction and 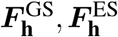 are the complex structure factors of the ground and excited states, respectively, for Miller index **h** = (*h, k, l*). This corresponds to Coppens’ “random-diffuse” model^18^. From Eq. (1), we see that the structure factors of the perturbed state can be expressed as a weighted average of the ground and excited-state structure factors:

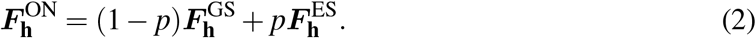

A central goal of perturbative crystallography experiments is to build a model of the excited state via refinement against excited-state structure factors. These must be estimated from the observed mixture of states. The ideal extrapolation method would start from Eq. (2), finding 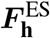 as

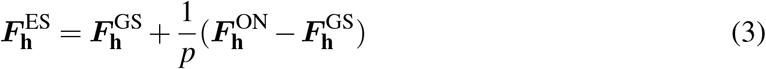

That is, extrapolation involves multiplying the vector difference 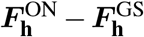 by the reciprocal of the excited-state fraction and then adding this amplified difference to the ground-state structure factor 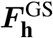 to recover the desired 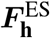 [Fig. 1(a)]. Of course, in practice one often does not know *p*. From here on, we will understand *p* as a “lens” with magnification 1*/p* that enhances the representation of excited-state density in the data, whether or not there is a specific excited state with that probability.

**FIG. 1.**
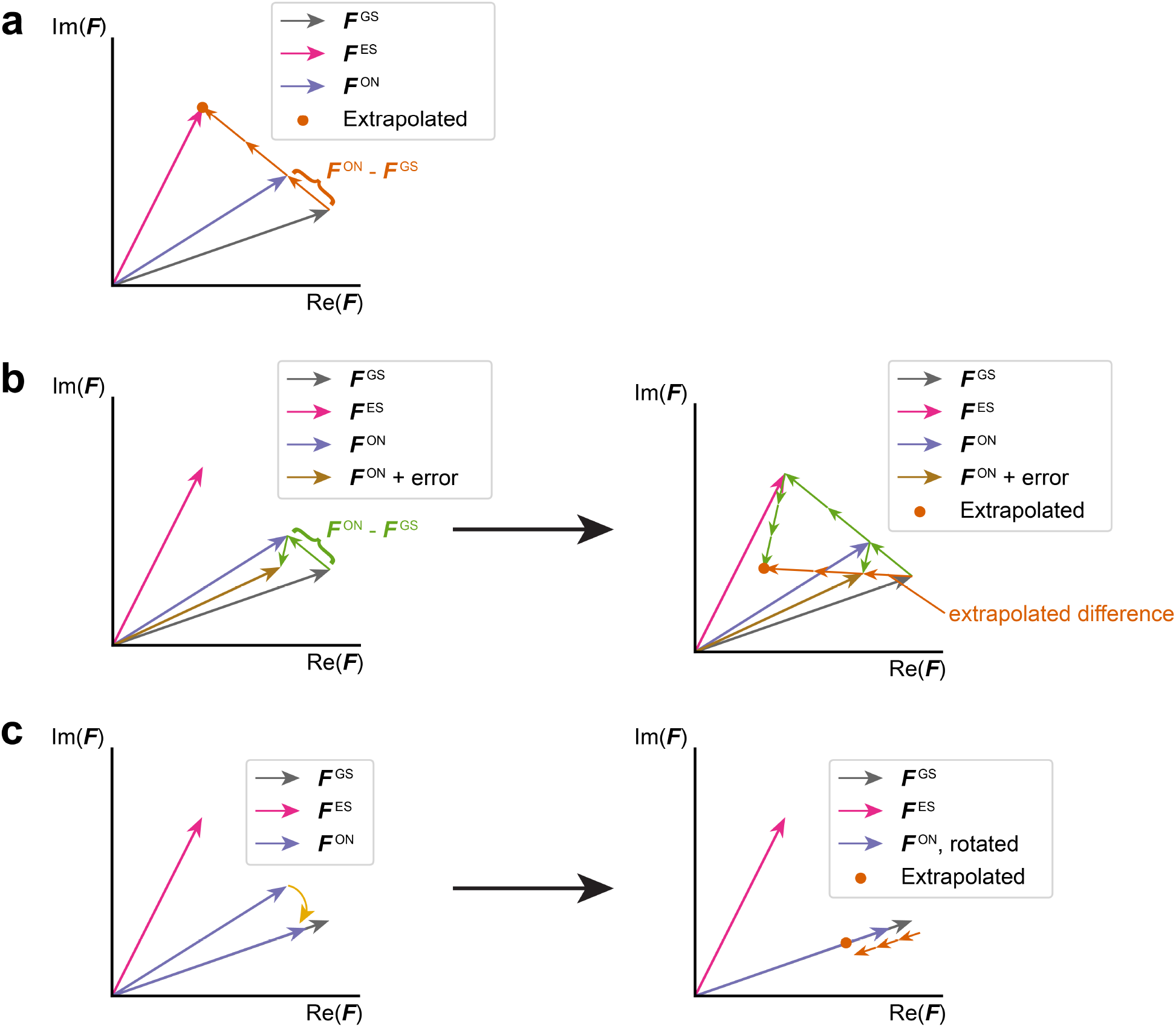
Theory and limitations of structure factor extrapolation. Structure factor extrapolation on a single reflection. The true ground-state and excited-state structure factors of this reflection are ***F***^GS^ and ***F***^ES^, respectively. The experimentally observed structure factor ***F***^ON^ has excited-state fraction 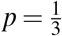. **(a)** In the complex plane, adding the extrapolated difference 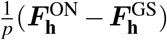 to 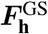 perfectly recovers the unobserved excited state 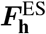. **(b)** Traditional extrapolation amplifies the true difference signal and any additive measurement errors by the same amount. Thus, even when errors are small relative to 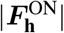, extrapolated structure factors can differ substantially from their true values. **(c)** Working with structure factor amplitudes in traditional extrapolation assumes that 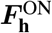 has the same phase as 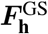. Traditional extrapolation is equivalent to rotating 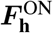 to have equal phase with 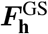 and then extrapolating the difference along the in-phase vectors. In the diagram, extrapolated difference vectors (orange) are offset from the collinear vectors for visibility. When phase differences are appreciable, extrapolated amplitudes become poor estimates of their true values.

However, crystallographic data reduction only estimates the amplitudes of complex structure factors (*F*_**h**_ = |***F***_**h**_|) and not their phases. As such, established methods for extrapolating excited-state structure factors^1,19^ apply the principle of Eq. (3) to structure factor *amplitudes*. The general form of this approach, which we refer to as traditional extrapolation, is:

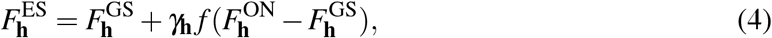

where the extrapolation factor *f* determines the extent of difference amplification and *γ*_**h**_, which can take various forms, is an optional weight on structure factor amplitude differences based on their relative magnitude and support under a Bayesian prior^19^. Taken together, the product of *γ*_**h**_ and *f* functions as 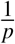.

There are several aspects of traditional structure factor extrapolation that limit its ability to produce excited-state structure factors that are suitable for refinement. First, naive extrapolation amplifies experimental noise just as much as it does the true difference in structure factor amplitudes, as evident in (Eq. 4). In any real experiment, the observed amplitude difference is a combination of the noise and the true difference in structure factor amplitudes. This can substantially impact the extrapolated amplitudes, even when the magnitude of errors is small relative to the amplitudes [Fig. 1(b)]. The weighting term *γ*_**h**_ in Eq. (4) is designed to address this issue by down-weighting reflections with high error estimates, thereby improving extrapolation in many cases^1,19^. Second, a limitation intrinsic to any kind of amplitude-based extrapolation is that structure factor amplitudes by themselves do not provide information on the sign of phase differences between the ground state and excited state. As a result, excited-state features in maps calculated by combining structure factor amplitude differences with ground-state phases are necessarily attenuated by half^20^. Indeed, in the absence of additional excited-state phase information, one typically extrapolates by treating 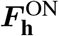 as collinear with 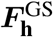 before amplifying their amplitude differences [Fig. 1 (c)]. Finally, traditional structure factor extrapolation can yield negative amplitudes—a problem with no generally agreed-upon remedy.

Here, we describe double-Wilson Extrapolator (hereafter, DW-Extrapolator), an algorithm that uses Bayesian inference to estimate extrapolated structure factor amplitudes. The approach relies on a statistical prior on complex structure factors (colloquially known as the double-Wilson distribution) that models the correlation between ground and excited-state structure factors^21^. From this prior, we can then infer 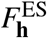 from the experimentally observed structure factor amplitudes of the ground and perturbed states.

DW-Extrapolator addresses some of the limitations of traditional extrapolation we described. First, the correlation imposed by the prior probabilistically constrains the magnitude of differences in structure factors between states, mitigating the effect of noisy structure factor amplitude differences. Second, since the double-Wilson model is a prior on the joint distribution of complex structure factors, we are able to perform inference with complex quantities, thereby explicitly modeling the distribution of the phase differences between datasets. This modeling cannot, by itself, overcome the fundamental ambiguity in the sign of the phase difference between ground and excited states, but optimally models the distribution of excited-state structure factors under the model assumptions—an improvement over more *ad hoc* approximations. Third, negative extrapolated structure factor amplitudes simply cannot occur in our framework since inference is performed with complex quantities. We implemented DW-Extrapolator in the rs-booster extension (https://github.com/rs-station/rs-booster) of reciprocalspaceship^22^.

After introducing the statistical theory and implementation of DW-Extrapolator, we evaluate the method’s performance on three benchmark perturbative crystallography experiments from the literature. The benchmarks consist of distinct model systems, perturbations, and crystallographic data processing software. Across benchmarks, DW-Extrapolator matches or exceeds the performance of conventional structure factor extrapolation calculations as implemented in Xtrapol8, a software platform for the application of such calculations.

## II. METHODS

### A. The Double-Wilson Model

Building on work by Read^23^, Hekstra *et al*. recently introduced the double-Wilson distribution as a statistical prior for modeling the correlation between closely related crystallographic datasets^21^. Although initially applied to scaling and merging reflection intensities from diffraction data, here we use the same statistical model for structure factor extrapolation. We now provide a description of the model.

Under the double-Wilson model, the normalized ground- and excited-state structure factors are jointly distributed according to a correlated complex normal distribution. Normalized structure factors are computed as 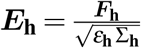, where *ε*_**h**_ is the reflection multiplicity due to crystal symmetry and Σ_**h**_ is a resolution-dependent scale factor accounting for systematic variation in scattering power across reciprocal space due to factors such as crystal mosaicity and molecular disorder^24,25^. For acentric reflections, structure factors can be expressed as:

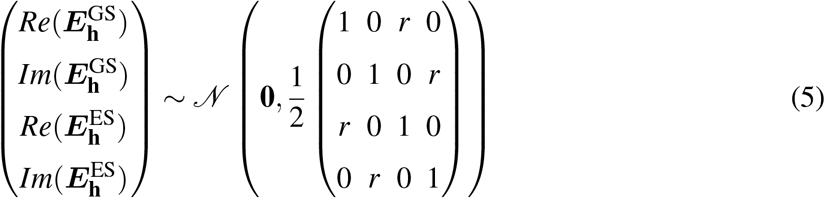

For centric reflections, which only have a real component, the distribution becomes:

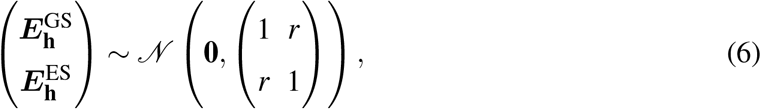

where the parameter *r* is the correlation between 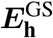 and 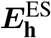.

### B. Bayesian Inference of Excited-State Structure Factor Amplitudes

Extrapolation requires estimating the true excited-state structure factor amplitudes 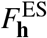 given the observed data. We now show how to estimate 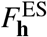 using a Bayesian approach with the double-Wilson prior. Like other approaches, our method closely resembles Eqs. (1)–(11) of Ursby and Bourgeois^26^ but avoids subsequent approximations. DW-Extrapolator takes as data two input datasets, one of the ground state (when the perturbation is “off”) and one of when the perturbation is turned “on”. The observed data can take two forms, depending on the processing software used:

1. Structure factor amplitudes 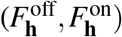 and associated error estimates 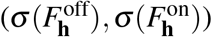
2. Integrated diffraction intensities 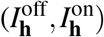 and associated error estimates 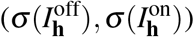

#### 1. Statistical Theory

In this section, we will first outline the inference framework and algorithm for the case where the provided data are structure factor amplitudes and their associated errors. DW-Extrapolator can also work with integrated intensities as data; the theory for this modified procedure is described in the Supplementary Text.

From a Bayesian perspective, we can treat the data as the observed values of random variables. We will denote the observed data with lowercase superscripts (off, on) and their associated random variables with uppercase superscripts (ES, GS, ON). Second, we treat—for now—our model parameters ***θ*** = (*r, p*) to be fixed quantities specified by the user. *r* is defined by Eqs. (5) and (6) for acentric and centric reflections, respectively. Importantly, we again note that *p* does not necessarily reflect the true fraction of excited-state molecules within the protein crystal. Indeed, as shown in the following analysis, *r* and *p* are hard to determine uniquely (or “identify”) without further structural modeling as they both affect the similarity between ground and excited-state structure factors^27^. As such, in the context of DW-Extrapolator, *p* should be understood only as a parameter tuning the extent of extrapolation.

To construct our Bayesian estimator for 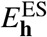, we are interested in the posterior distribution of 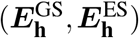 given the observed data and the statistical priors described above:

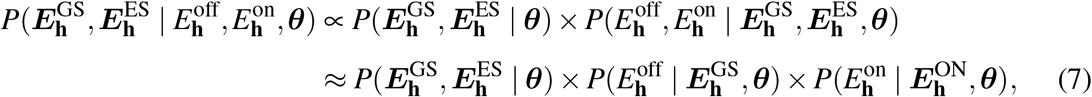

where 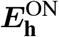 is defined according to (2). We approximate the measurement errors of the “off” and “on” datasets as independent, allowing us to treat the observed structure factor amplitudes as conditionally independent given their true values. For experimental structure factor amplitudes obtained by French-Wilson scaling^24^ (e.g., in XDS^28^) or by scaling in Careless^29^, the likelihood functions are given by the truncated normal distribution (see Supplementary Text IA). To account for the proportionality in the first step, we use self-normalized weights in the importance sampling procedure described below. We aim to recover the posterior mean as the estimator for 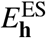:

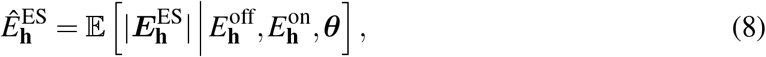

and the posterior variance as the estimator for the associated error:

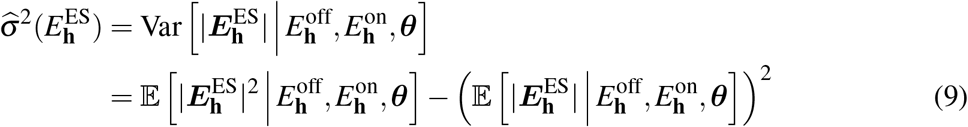

#### 2. Monte Carlo Integration Algorithm

To approximate the expectation values in Eqs. (8) and (9), DW-Extrapolator uses importance sampling to perform Monte Carlo integration. Users provide data for the “off” and “on” datasets in the form of .mtz files. Additionally, users specify the parameter values ***θ*** = (*r, p*).

The algorithm proceeds as follows:

1. Sample *n* = 10^6^ i.i.d. multivariate normal vectors as **x**_*i*,4×1_ ∼ 𝒩 (**0, I**_4×4_), collecting results in a matrix **X**_4×*n*_. These are the samples for acentric reflections, with sample index *i*.
2. Sample *n* = 10^6^ i.i.d. multivariate normal vectors as **y**_*i*,2×1_ ∼ 𝒩 (**0, I**_2×2_), collecting results in a matrix **Y**_2×*n*_. These are the samples for centric reflections.
3. Transform **X**_4×*n*_ and **Y**_2×*n*_ so that the multivariate normals covary as the double-Wilson distribution (covariance matrices ***C*** for acentrics, ***C***^***′***^ for acentrics to match Eqs. (5) and (6), respectively). Specifically, create 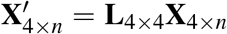 and 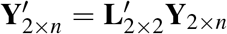, where ***C*** = **LL**^*T*^ and ***C***^***′***^ = **L**^′^**L**^′*T*^. We obtain **L** and **L**^′^ by Cholesky decomposition. Each column *i* in 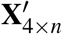 and 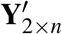 is then a sample from the Double-Wilson prior for acentric and centric reflections, respectively:

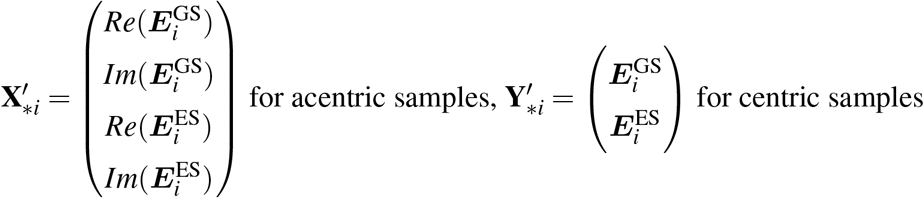
4. For each sample *i*, compute

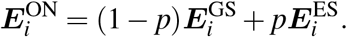

Then apply a single scalar correction factor to all samples so that the sample median of |***E***^ON^| matches that of |***E***^GS^|. This correction is necessary because affine combinations of normalized structure factors are not normalized themselves, while normalization is assumed during preprocessing when computing the scale factors Σ_**h**_. We use the median of |***E***^ON^| as a robust alternative to the mean or root-mean-square when matching magnitudes.**Then, for each Miller index h:**
5. Follow Eq. (7) to compute joint likelihood weights *w*_**h**,*i*_ for each sample 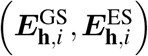 given the observed structure factor amplitudes:

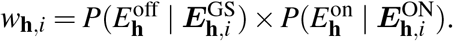

Normalize weights as 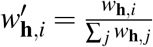
6. Approximate the expectation value in Eq. (8) as: 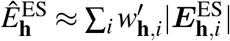Associated error estimates from Eq. (9) follow: 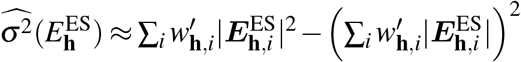
7. Place structure factors and their associated errors on their original scales according to:

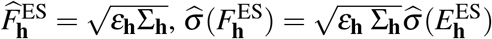

In what follows, we will refer to 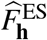 as *F*_extr,**h**_ where that improves clarity. Steps (5) and (7) rely on interconversion between normalized structure factor amplitudes and observed ones. To do so, we use reciprocalspaceship^22^ to calculate multiplicities *ε*_**h**_ and estimates of scales Σ_**h**_ = ⟨*I*_**h**_⟩, with the latter indicating the average intensity near reciprocal lattice point **h**.

Having described the inference procedure when working with structure factor amplitude data, we note that many crystallographic processing methods (e.g., DIALS^30^ and Aimless^31^) estimate diffraction intensities. In these cases, DW-Extrapolator takes the integrated diffraction intensities as data and uses a Normal likelihood function given 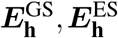, ***θ*** (see Supplementary Text IB).

### C. Maximum Likelihood Estimation of Model Parameters

Recall that DW-Extrapolator requires users to specify the parameters ***θ*** = (*r, p*). Alternatively, maximum likelihood estimation (MLE) can provide parameter estimates informed by the data and the statistical model. When the data are structure factor amplitudes, the likelihood function for our parameters ***θ*** given the observed data is:

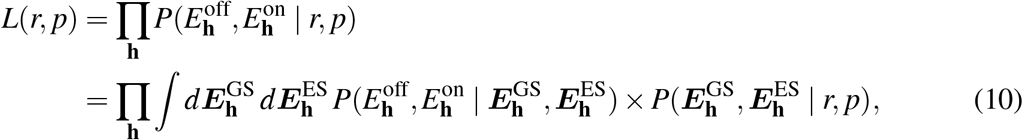

with negative log likelihood (NLL) function following as:

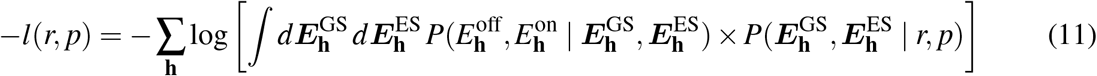

When the provided data are instead integrated intensities, the per-reflection likelihood functions in Eqs. (10) and (11) are replaced by their intensity equivalents (Supplementary Text IB).

As before, we use importance sampling to approximate the integrals in Eq. (11). Then, using the scipy.optimize package^32^ and Eq. (11) as the objective function, we can infer maximum-likelihood estimates 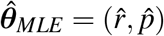.

### D. Test Cases

To assess the performance of DW-Extrapolator, we selected three examples of perturbative crystallography datasets from the literature. Across these benchmarks, we sought to sample a representative range of protein types, perturbations, and data processing software. Those details are summarized in Table I.

**TABLE I.** Summary of Benchmark Experiments.

| Protein | Perturbation | PDB IDs<br>(Ground State, Excited State) | Reflections<br>Dataset | Data Processing<br>Software |
| --- | --- | --- | --- | --- |
| Photoactive Yellow Protein | Chromophore excitation | 1TS0 (A chain),<br>1TS0 (B chain)<sup>33</sup> | Diffraction images<sup>29</sup><br>collected for Dalton *et al.* | Careless<sup>29</sup> |
| BAZ2B | Ligand binding | From fragment screening dataset<br>in Pearce *et al.*<sup>15</sup> | Same as PDBs | Aimless<sup>31</sup> |
| mEos4b | Chromophore excitation | 6GP0, 6GP1<sup>34</sup> | Same as PDBs | XDS<sup>28</sup> |

Each test case has published ground and excited-state models that we will treat as the ground truth. The ground-state models represent the true unperturbed state given accurate modeling choices. However, it is a stronger assumption that the excited-state models are accurate. Still, as our test cases undergo biochemically well-characterized perturbations with clear structural consequences (e.g., a chromophore flip or a ligand-binding event), the resulting structural models are likely to represent the excited states accurately.

### E. Running Xtrapol8

As a point of comparison for DW-Extrapolator, we used Xtrapol8 (version 1.2.9)^19^. Xtrapol8 offers nine different strategies for traditional extrapolation that differ in two ways: first, in their weighting schemes; second, in whether they extrapolate from observed or calculated ground-state structure factors.

For each test case, we ran Xtrapol8 first in its “fast-and-furious” mode, which performs a wide scan over potential occupancy values (i.e., excited-state fractions) using a subset of the nine available strategies. The output of this initial “fast-and-furious” run also includes an estimate of the excited-state occupancy. We then ran Xtrapol8 in its “calm-and-curious” mode with default settings. For these runs, we used all nine available extrapolation methods and tried a narrow grid of 25 evenly-spaced occupancy values around the estimated occupancy.

### F. Extrapolation Performance Metrics

To compare the performances of DW-Extrapolator and Xtrapol8, we used three metrics, described below. When making these comparisons in the below analysis, we consider for each method both how performance varies with model parameters and also the single parameter combination that produced the best metric.

#### 1. Difference Map Real-Space Correlation Coefficients

First, we sought to evaluate the effect of extrapolation on difference map quality. To do so, we first calculated the ground-truth difference map 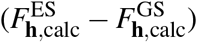 using calculated structure factors from published structures. Then, we compared this to the extrapolated difference density map 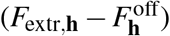 by computing the real-space correlation coefficient (RSCC) between the two maps (recall 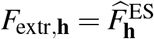). To evaluate map quality near expected structural changes, RSCC was computed in a 10 Å radius around the chromophore for the PYP and mEos4b datasets. For the BAZ2B drug-fragment screen, RSCC was computed in a 2 Å mask around each atom of the bound fragment. Real-space difference maps were computed using custom scripts based on gemmi^35^ and reciprocalspaceship^22^. Model structure factors were calculated using phenix^36^. To down-weight the contribution of particularly noisy reflections, we computed structure factor amplitude differences according to:

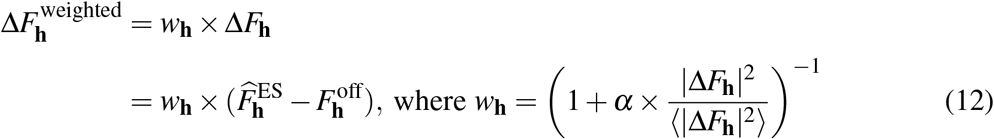

and the brackets indicate an average across all Miller indices in the dataset. This weighting scheme is based on the one introduced by Ren *et al*.^37^ but excludes the Bayesian weight term since our extrapolation procedure already has a Bayesian structure. For our difference maps, we chose *α* = 0.05^4^, and this weighting scheme improved difference map signal strength [Fig. S1(a)]. To ensure consistency of comparison, the calculated difference maps were cut to the same resolution as the extrapolated difference maps.

#### 2. R-Factors

Second, we used crystallographic *R*-factors to measure the agreement between the extrapolated structure factor amplitudes and the published models. We computed *R*-factors for the extrapolated structure factor amplitudes against the published ground-state (*R*_GS_) and excited state (*R*_ES_) structures. We then also computed their difference Δ*R* = *R*_ES_ − *R*_GS_. Generally, extrapolation leads to larger *R*-factors due to amplification of measurement errors. A small—or, better yet, negative— Δ*R* signifies that the extrapolated structure factor amplitudes are a better fit for the excited-state model than for the ground-state model, suggesting that extrapolation successfully biases the structure factor amplitudes towards capturing features of the excited state. *R*-factors were computed using a custom script based on cctbx^38^.

#### 3. 2*mF*_*extr*_ − 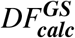 Map RSCCs

Finally, the ultimate goal of structure factor extrapolation is to refine structural models of the excited state. To get a sense of how well extrapolated structure factor refinement might proceed from a ground-state model, we computed 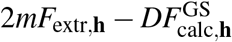 maps using extrapolated structure factor amplitudes and the published ground-state structures based on our analysis and based on Xtrapol8. These weighted difference maps are commonly used in refinement as an unbiased representation of the electron density supported by both the experimental data and the current model; therefore, a useful 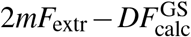 map should closely resemble the electron density of the excited state. As such, we computed RSCCs between these maps and 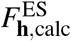 electron density calculated from the excited-state models. To focus on the relevant structural changes, RSCC was computed in a 2 Å mask around each atom of the chromophore (PYP and mEos4b datasets) or ligand (BAZ2B dataset) molecule. We generated 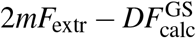 maps using phenix^36^ and computed RSCCs with custom scripts based on gemmi^35^ and reciprocalspaceship^22^. As before, to ensure consistency of comparison, calculated excited-state structure factors were cut to the same resolution as the 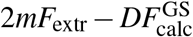 maps.

### G. DW-Extrapolator Usage

DW-Extrapolator is implemented in the dw_extrapolate command in the rs-booster package that supplements the reciprocalspaceship Python library^22^. The minimal arguments that users should supply are:

- `off-mtz:` .mtz file for unperturbed data
- `on-mtz:` .mtz file for perturbed data
- `r`: estimated correlation between ground and excited states on [0, 1]
- `p`: excited-state fraction on [0, 1]

For data merged in Careless^29^, which specifies the parameters of the truncated normal likelihood function for each observed structure factor amplitude, no additional arguments are needed. For data prepared by other software, we first check if their outputs contain integrated intensities and associated errors, in which case DW-Extrapolator should be run with the `–use_intensities` argument. The final use-case is for structure factor amplitudes that have been truncated or French-Wilson scaled^24^ but not explicitly parameterized in terms of a truncated normal distribution (see Supplementary Text IA for a description of how this case is treated). For these data, DW-Extrapolator should be run with the `–use_structure_factors` argument [Fig. 2]. For fastest performance, we recommend running DW-Extrapolator across multiple cores to allow for parallel processing.

**FIG. 2.**
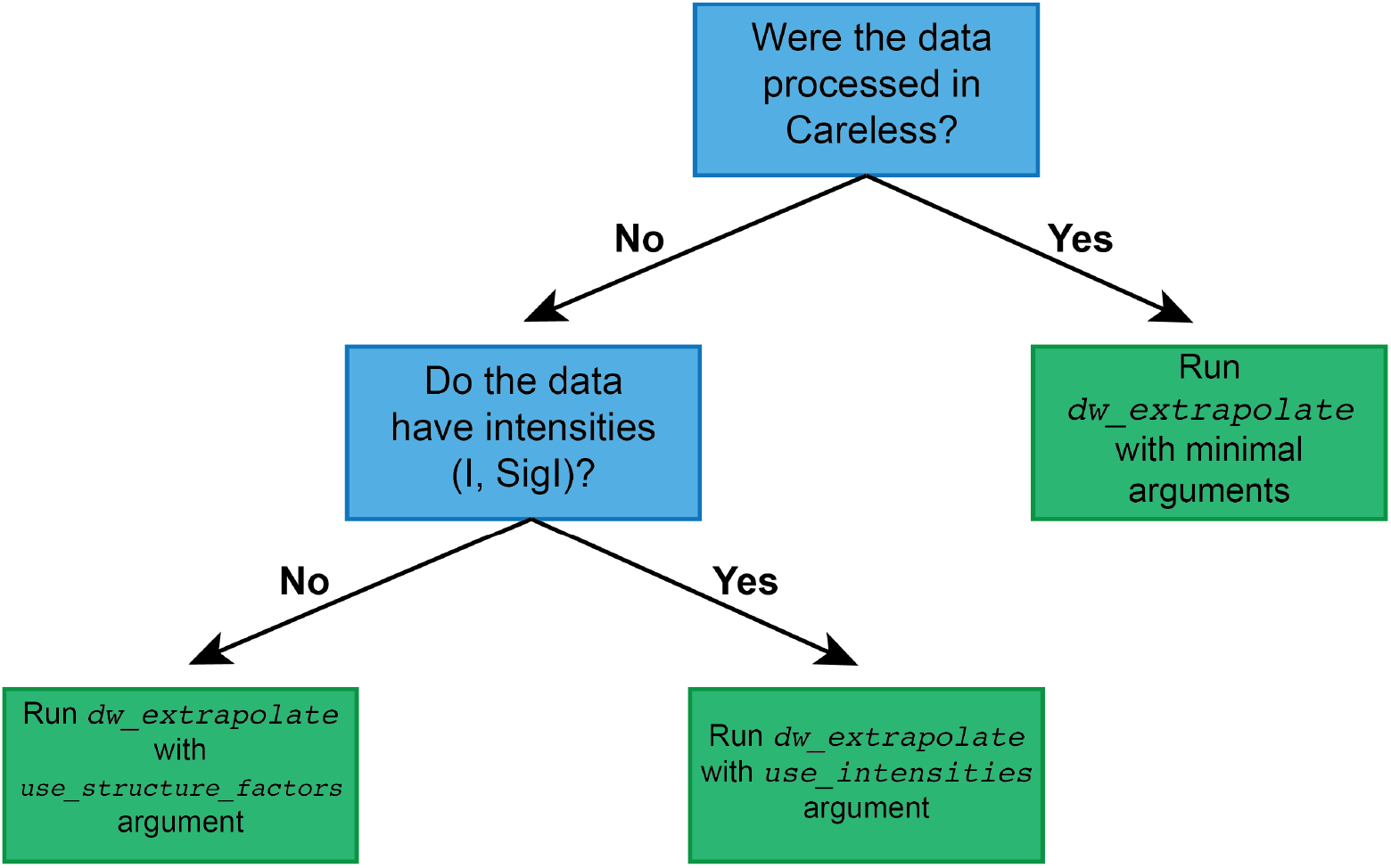
Decision tree for selecting the inference protocol to use in DW-Extrapolator. Crystallographic data processing software differ in the physical and statistical methods used to generate structure factor amplitudes from diffraction intensities. DW-Extrapolator can work with data from a comprehensive range of processing algorithms.

To provide users an initial set of parameters ***θ*** = (*r, p*) to try, the `dw_extrapolate` command contains a `–default_scan` option, which fixes *r* = 0.9 and then scans over ten values of *p* ∈ {0.05, 0.1,…0.45, 0.5}. For each parameter combination, the method produces extrapolated structure factor amplitudes and outputs the associated NLL. We recommend using the parameters that achieve the lowest NLL in this 1D scan as a starting point. The default choice of *r* = 0.9 has worked well in our experience for datasets with strongly correlated ground and excited states. If users have reason to believe that their ground and excited states are significantly more or less similar, however, they are able to specify a different *r* value.

In addition to this 1D parameter scan, the command `mle_dw_extrapolate` on rs-booster finds the MLE estimate for (*r, p*) across all of the two-dimensional parameter space.

## III. RESULTS

Given two diffraction datasets corresponding to a protein’s “off” state and its perturbed “on” state, DW-Extrapolator performs Bayesian inference to estimate the structure factor amplitudes of the isolated excited state 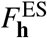 for all common Miller indices **h** between the two datasets. DW-Extrapolator’s inference procedure requires parameters ***θ*** = (*r, p*). *r* is the assumed correlation between the corresponding components of the complex structure factors 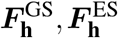. Higher values of *r* tend to produce estimates 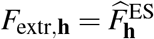 that are more similar to the observed ground-state structure factor amplitudes. *p* is the assumed excited-state fraction of the sample; however, estimates of *p* should not be directly interpreted as the true excited-state fraction due to the unidentifiability of *r* and *p* as well as the lack of phase information for the excited state^20^. Larger values of *p* produce estimates of the excited-state structure factor amplitudes closer to the perturbed data. Low values of *r* and *p* provide flexibility for inferring excited-state amplitudes that may differ greatly from the observed data.

Below, we assess DW-Extrapolator’s performance on three perturbative crystallography benchmark datasets from the literature, comparing its output to traditional approaches as implemented in Xtrapol8. We will take three approaches to selecting *r* and *p*: (a) use of a 2D-grid for sets *r* = 1 − 2^−*n*^, with *n* = 0, 1, … 11, and *p* = 2^−*m*^, with *m* = 0, 1, … 8; (b) maximum likelihood estimation of (*r, p*); and (c) a practical 1D scan as an alternative to a full scan of (*r, p*). After establishing how extrapolation results vary across parameter space, we then evaluate whether our likelihood-based parameter estimates perform well across key metrics. Finally, we will assess DW-Extrapolator’s performance across parameter space and provide recommendations for practical use.

For our three examples, we find that DW-Extrapolator successfully infers features of the excited states while mitigating the extent to which extrapolation amplifies errors. Although our metrics are not perfectly correlated, our inference methods identify parameter combinations that perform well in all three evaluation criteria. These results support the validity of DW-Extrapolator’s statistical framework and its utility for a range of perturbative crystallography contexts.

We also compare DW-Extrapolator’s performance to the nine distinct extrapolation strategies implemented in Xtrapol8, a state-of-the-art software interface for structure factor extrapolation. For each strategy, we scan a range of recommended values of the estimated excited-state occupancy. In Figs. 3-5, we primarily compare to whichever of these nine strategies performs best based on each criterion. We find that the “k-weighting” scheme based on work by Ren *et al*.^37^ performs consistently well across benchmarks and evaluation metrics. We also compare our parameter recommendations to Xtrapol8’s recommended weighting strategy and occupancy values, which are computed using difference map metrics^19^.

**FIG. 3.**
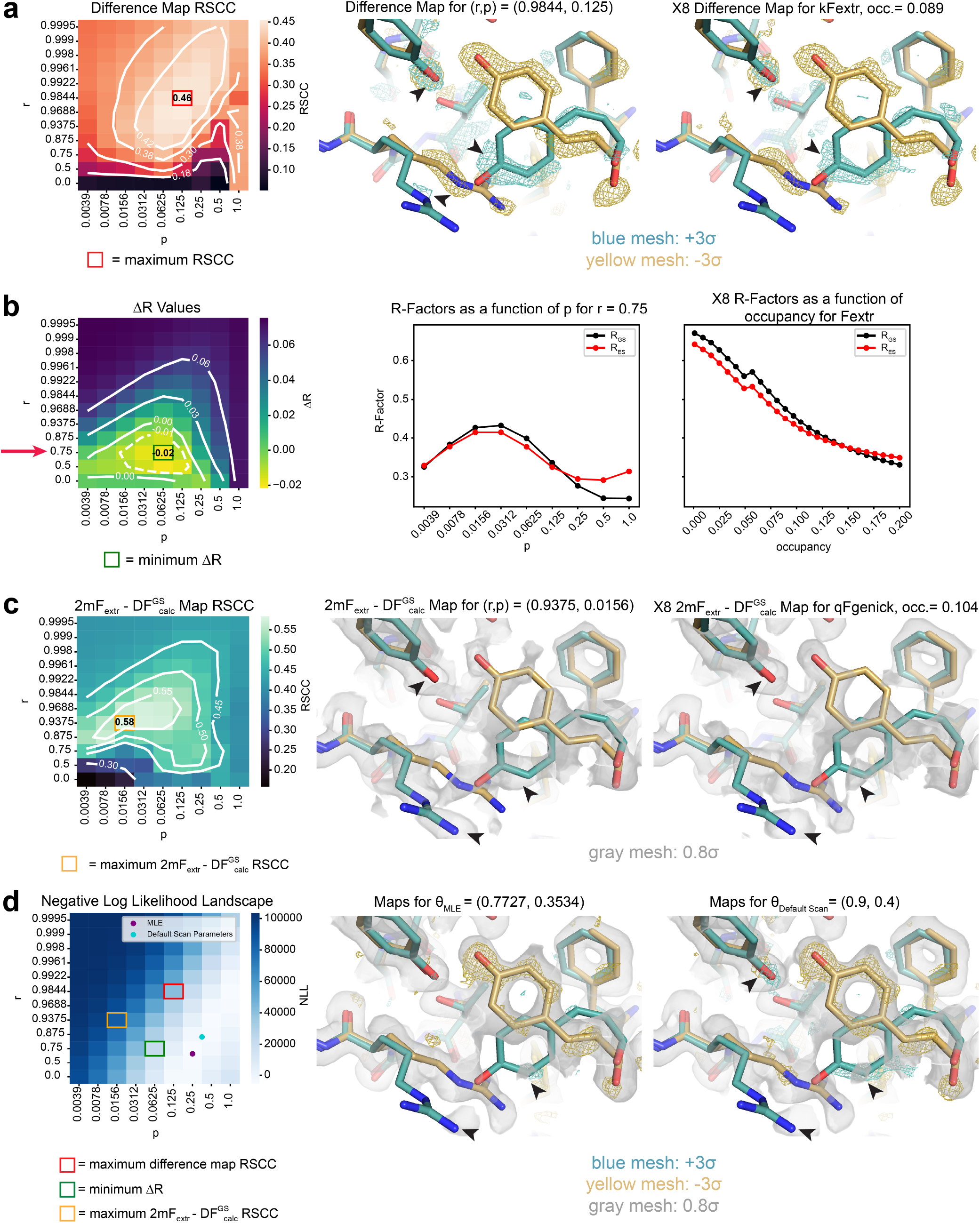
DW-Extrapolator of a PYP chromophore excitation dataset. **(a)** Difference Map RSCC values within 10 Å of the chromophore across a grid of (*r, p*). Isocontour lines are drawn in white. The molecular models on the right show the PYP chromophore site of the ground (yellow) and excited (teal) states, as well as the difference maps (meshes) that produced the highest RSCCs in the grid searches for DW-Extrapolator and Xtrapol8 (X8). The difference density is contoured to ±3*σ*. Arrowheads indicate positive difference density corresponding to features of the excited state. **(b)** Δ*R*-factor values across the (*r, p*) grid with isocontour lines are drawn in white. The plot in the middle panel fixes *r* = 0.75 and visualizes how *R*-factors change as a function of *p*. The plot on the right selects X8’s Fextr weighting strategy, which achieved the smallest Δ*R* in the occupancy grid, and plots *R*-factors as a function of occupancy. **(c)** 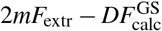 map RSCC values across (*r, p*) computed in a 2 Å mask around the *p*-coumaric acid chromophore (here, 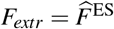). Isocontour lines are drawn in white. The molecular model shows the PYP chromophore with electron density from the 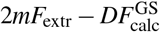 maps that produced the highest RSCCs in the grid searches for DW-Extrapolator and X8. The electron density is contoured to 0.8*σ*. Arrowheads indicate map density corresponding to features of the excited state. **(d)** Negative Log Likelihood landscape of ***θ*** = (*r, p*) under the double-Wilson model given the experimental data. The maximum likelihood estimate 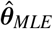 and default scan parameters ***θ***_Default Scan_ suggested by DW-Extrapolator are plotted on the heatmap. The molecular models show the PYP chromophore with the difference (teal/yellow meshes) and 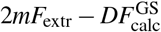 (gray) maps produced by structure factor amplitudes extrapolated with these recommended parameters. The maps are contoured as before. Arrowheads indicate either positive difference density or 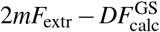 density corresponding to features of the excited state.

### A. Photoactive Yellow Protein

To benchmark DW-Extrapolator, we start with data measured on crystals of Photoactive Yellow Protein (PYP), a well-characterized photoreceptor that responds to blue light stimulation^39^. In the first TR-X studies, researchers solved structures before and after exposing a PYP crystal to blue light, showing that the PYP photocycle initiated with the *trans* to *cis* isomerization of its *p*-coumaric acid chromophore^1,33^. We apply DW-Extrapolator to a polychromatic dataset of 20 images of PYP in its “dark” ground state and 20 images 2 ms after photoexcitation, at which point the chromophore is in its perturbed state. These data were scaled and merged in Careless^29^. We find that DW-Extrapolator successfully captures density corresponding to the PYP chromophore’s *cis* state [Fig. 3(a)]. We now quantify DW-Extrapolator’s success with several metrics.

To measure how well DW-Extrapolator can detect difference signal resulting from chromophore isomerization, we compare difference maps computed with structure factor amplitudes produced by DW-Extrapolator to a ground-truth difference map using a real-space correlation coefficient (RSCC) as described in Methods II F 1. The best RSCC of a DW-Extrapolator difference map (0.46), swept across our grid of *r* and *p*, is higher than that of an unextrapolated difference map calculated from the observed “dark” and 2 ms datasets [Table II]. Indeed, difference density for the chromophore’s *cis* conformation is strongly present in the DW-Extrapolator difference map [Fig. 3(a)]. Taken together, these two observations indicate that DW-Extrapolator enhances difference map quality. To investigate how these maps compare to those produced by traditional extrapolation, we apply Xtrapol8 to this dataset. The “k-weighting” strategy produces the highest quality difference maps, with a maximum achieved RSCC of 0.42 [Fig. S2(a)]. These results indicate that DW-Extrapolator can detect difference density more strongly than difference maps across various weighting strategies in traditional extrapolation.

**TABLE II.** Difference Map RSCC Values.

| Protein | Unextrapolated | DW-Extrapolator<br>2D Grid Max. | Xtrapol8<br>Grid Max. | DW-Extrapolator MLE | DW-Extrapolator<br>Default Scan | Xtrapol8<br>Recommended<br>Occupancy |
| --- | --- | --- | --- | --- | --- | --- |
| Photoactive Yellow<br>Protein (PYP) | 0.38 | 0.46 | 0.42 | 0.18 | 0.30 | 0.36 |
| BAZ2B | 0.70 | 0.78 | 0.78 | 0.70 | 0.72 | 0.74 |
| mEos4b | 0.48 | 0.60 | 0.47 | 0.57 | 0.57 | 0.45 |

To determine how well excited-state maps from DW-Extrapolator aid structural refinement, we next assess the quality of DW-Extrapolator’s excited-state maps. One potential quality metric is the crystallographic *R*-factor of the extrapolated map against the ground-truth model, where a smaller *R*-factor indicates better agreement. However, since traditional extrapolation amplifies errors, the resulting maps are often overall of poorer quality and have higher *R*-factors against the ground-state model, even if they capture key features of excited states. Therefore, to measure how well the excited-state map captures features strictly corresponding to excited-state structures and not error minimization, we instead use the *R*-factor difference Δ*R* [see Methods II F 2]. A negative value of Δ*R* indicates that the extrapolated structure factor amplitudes are a better fit for the excited-state ground-truth model than the ground-state model.

We find that for PYP, DW-Extrapolator produces structure factor amplitudes with Δ*R <* 0. For a fixed correlation parameter *r*, tuning the excited-state fraction *p* can shift this bias [Fig. 3(b)]. We rationalize this behavior as follows. For high *p*, which corresponds to the assumption that the excited state dominates the observed data, the estimated *F*_extr,**h**_ is underextrapolated and overrepresents the contribution from the ground state. This results in the expected *R*_GS_ *< R*_ES_. Then, for intermediate values of *p* that likely approach the true excited-state fraction, *R*_ES_ *< R*_GS_, indicating that extrapolation properly infers features of the excited-state structure. For small *p, R*_GS_ *< R*_ES_ once again. In this regime, the method “overextrapolates,” as it assumes that the observed mixture of states provides very little information on the excited state. The likelihood terms of the posterior given in (7) then become small, and the prior term dominates. Since the double-Wilson prior enforces similarity with the ground state, it follows that the extrapolated structure factor amplitudes will again resemble the ground state.

We find that the *R*-factors for DW-Extrapolator structure factor amplitudes are much lower than those computed by Xtrapol8 when also using observed, rather than calculated, ground-state structure factor amplitudes as the reference. For PYP, the *R*-factors that achieve the biggest Δ*R* gap with DW-Extrapolator are around 10 percentage points smaller than those for Xtrapol8 [Table III]. Indeed, *R*-factors for structure factor amplitudes from DW-Extrapolator tend to be lower across excited-state fractions [Fig. S1(b)]. For Xtrapol8, we observe a monotonic increase in *R*-factors with increasing extrapolation, which is consistent with amplification of errors during traditional extrapolation [Fig. 3(b)]. We believe that DW-Extrapolator achieves much lower *R*-factors because the prior’s correlation structure effectively suppresses large errors. Since *R*-factors report on the average deviation *F*_obs_ − *F*_calc_ across reflections, and *F*_calc_ is kept fixed in this comparison, we conclude that the inferred “observed” excited-state structure factor amplitudes produced by DW-Extrapolator are therefore more accurate.

**TABLE III.** (*R*_GS_,*R*_ES_) *R*-factor Values.

| Protein | DW-Extrapolator<br>2D Grid Min. $\Delta R$ | Xtrapol8<br>Grid Min. $\Delta R$ | DW-Extrapolator MLE | DW-Extrapolator<br>Default Scan | Xtrapol8<br>Recommended<br>Occupancy |
| --- | --- | --- | --- | --- | --- |
| Photoactive Yellow<br>Protein (PYP) | (0.40, 0.38) | (0.57, 0.53) | (0.26, 0.29) | (0.25, 0.30) | (0.61, 0.58) |
| BAZ2B | (0.29, 0.28) | (0.47, 0.46) | (0.32, 0.31) | (0.27, 0.26) | (0.43, 0.42) |
| mEos4b | (0.41, 0.38) | (0.61, 0.54) | (0.26, 0.26) | (0.24, 0.25) | (0.48, 0.43) |

Typically, excited-state models are manually refined against weighted difference maps. To determine whether DW-Extrapolator produces high-quality 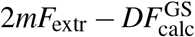 excited-state maps, we measure the RSCC of these maps to corresponding ground-truth excited-state models [see Methods II F 3]. DW-Extrapolator’s best 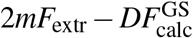 map RSCC is greater than that achieved by the unextrapolated map and the methods implemented in Xtrapol8 [Table IV]. This best DW-Extrapolator 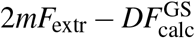 map displays density supporting features of the PYP active site’s excited state, such as the alternate conformations of the *p*-coumaric acid ring and the adjacent R52 residue [Fig. 3(c)].

**TABLE IV.** 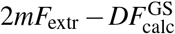 Map RSCC Values.

| Protein | Unextrapolated | DW-Extrapolator<br>2D Grid Max. | Xtrapol8<br>Grid Max. | DW-Extrapolator MLE | DW-Extrapolator<br>Default Scan | Xtrapol8<br>Recommended<br>Occupancy |
| --- | --- | --- | --- | --- | --- | --- |
| Photoactive Yellow<br>Protein (PYP) | 0.40 | 0.58 | 0.55 | 0.48 | 0.44 | 0.49 |
| BAZ2B | 0.83 | 0.90 | 0.87 | 0.84 | 0.86 | 0.69 |
| mEos4b | 0.87 | 0.89 | 0.89 | 0.89 | 0.89 | 0.75 |

#### Finding suitable parameters (r, p)

In the absence of structural models of the excited state to compare to, it can be unclear how to choose *r* and *p*. In principle, our formalism permits direct optimization of the likelihood of the two parameters. To examine this, we implemented a negative log-likelihood (NLL) calculation in `dw_extrapolate` that is returned along with each output .mtz. Further, `mle_dw_extrapolate` performs maximum likelihood estimation (MLE) to infer plausible parameters 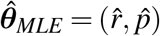. Plotting the NLL over the 2D (*r, p*) grid for the PYP dataset reveals a “valley” in parameter space along which the NLL is minimized [Fig. 3(d)]. The parameter values that have the best performance as measured by difference map RSCC and Δ*R* lie near—although often slightly above— this valley, confirming that the NLL calculation can inform the choice of *r* and *p*.

For the PYP example, structure factors extrapolated with 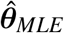 have excited-state model *R*-factor *R*_ES_ = 0.29 and produce a 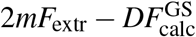 map with density for the excited-state chromophore [Fig. 3(d); Tables III, IV]. Therefore, extrapolation with MLE parameters provides a good starting point for structure refinement. The difference map produced by these MLE structure factor amplitudes has a lower RSCC than the unextrapolated difference map [Table II]. Visualizing the map reveals strong negative difference density that would be helpful during refinement; we attribute the map’s low RSCC to its comparatively weak positive difference density [Fig. 3(d)].

However, the presence of a valley of parameters with similar likelihood means that only the combination of parameters (*r, p*) can be well estimated. Empirically, we find in each test case [Fig. S1(d), S3(d), S5(d)] that these NLL valleys are well approximated by contours defined by:

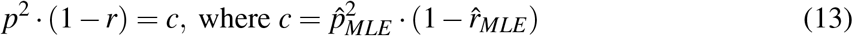

Accordingly, a practical approach is to fix *r* at a reasonable prior value. In practice, correlations between related datasets are often quite high (>0.8)^21^, and a 1D scan of *p* will suffice. Current approaches ignoring expected correlations are equivalent to choosing *r* = 0. Choosing a value of *r* that is too high will skew the excited state to look like the ground state—a conservative bias. As a reasonable estimate, DW-Extrapolator’s default scan option uses *r* = 0.9 but also gives users the option to specify *r*.

Within this 1D scan for the PYP dataset, *p* = 0.4 gives the highest likelihood. The resulting structure factor amplitudes extrapolated with ***θ***_Default Scan_ = (0.9, 0.4) have comparable performance to those produced with 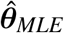 as measured by *R*-factors and 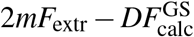 RSCC, while having a higher difference map RSCC [Tables II-IV]. Visualizing the maps for these structure factor amplitudes shows strong positive and negative difference density, as well as 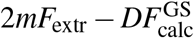 density for the chromophore’s excited-state conformation [Fig. 3(d)]. This comparison suggests that a 1D scan with a reasonable *r*-value performs as well as the maximum-likelihood estimate optimized over the 2D parameter space.

As a practical use comparison, we take Xtrapol8’s recommendation of “q-weighting” with occupancy of 0.049. These extrapolated structure factor amplitudes yield slightly higher difference and 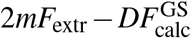 map RSCCs than achieved by DW-Extrapolator’s likelihood-based parameter estimates [Tables II, IV; Fig. S2(d)]. Yet, the *R*-factors associated with Xtrapol8’s recommended structure factor amplitudes are around 30 percentage points higher [Table III].

### B. BAZ2B Ligand Screening

For our second example, we apply DW-Extrapolator to a drug-fragment screening dataset for the bromodomain BAZ2B^15,40^. The ground state in this experiment is the protein in its *apo* form, and the excited state is the *holo* protein bound to a drug fragment. Note that in the *apo* protein, the ligand binding site is occupied by a molecule from the crystallization solution, which Pearce *et al*. chose to model as a *1,2*-ethanediol molecule. Additionally, when De Zitter *et al*. used this dataset to validate Xtrapol8, they set a high-resolution limit of 1.85 Å as the *apo* and *holo* datasets displayed non-isomorphism towards high resolution^19^. For consistency of comparison, we use the same resolution limit for our application of DW-Extrapolator. Finally, the reflection data from these experiments were processed using Aimless^31^, so DW-Extrapolator was run with the `–use_intensities` argument.

To assess whether DW-Extrapolator can capture signal for small-molecule binding events, we calculate difference maps using the extrapolated structure factor amplitudes. Even though the unextrapolated difference map already shows strong ligand density, DW-Extrapolator improves the difference map quality (as measured by RSCC with the calculated ground-truth difference map) such that the electron density better follows the actual shape of the ligand molecule [Fig. S4(a), Fig. 4(a)]. For reference, traditional extrapolation as implemented in Xtrapol8 achieves a maximum difference map RSCC equal to the best in our grid of (*r, p*) parameters for DW-Extrapolator [Table II].

**FIG. 4.**
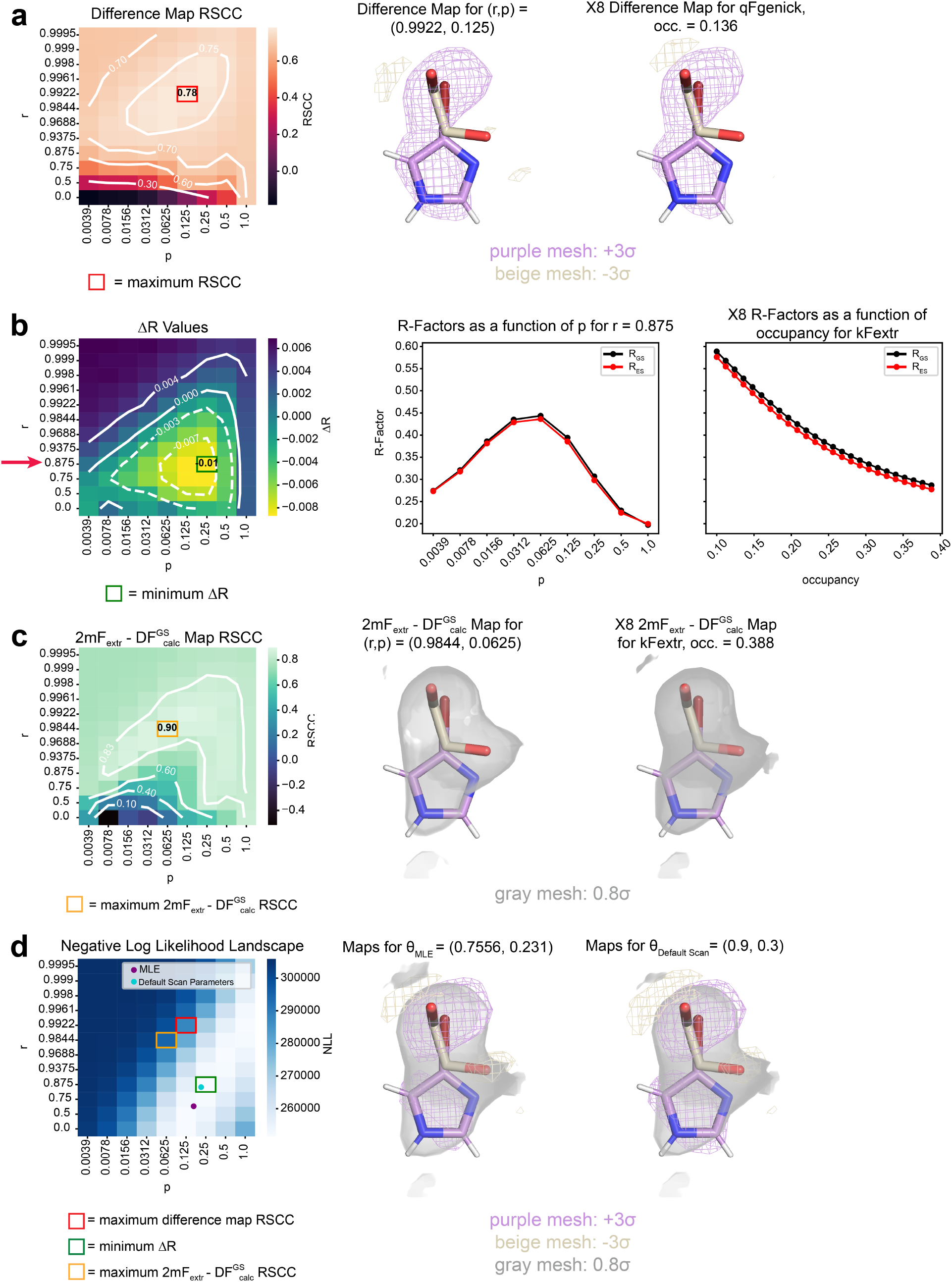
DW-Extrapolator results for the BAZ2B fragment screening dataset. **(a)** Difference Map RSCC values in a 2 Å mask around the ligand across a grid of (*r, p*). Isocontour lines are drawn in white. The BAZ2B ligand binding site is visualized in its *apo* (beige) and *holo* (purple) states. The difference density for the maps with the best RSCCs in the grid searches for DW-Extrapolator and Xtrapol8 (X8) is contoured to ±3*σ*. **(b)** Δ*R*-factor values across the (*r, p*) grid with isocontour lines drawn in white. The center plot fixes *r* = 0.875 and visualizes how *R*-factors change as a function of *p* for structure factor amplitudes produced by DW-Extrapolator. The plot on the right visualizes how *R*-factors change as a function of occupancy for the kFextr extrapolation strategy in X8. **(c)** 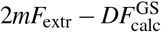 map RSCC values across (*r, p*) calculated for a 2 Å mask around the ligand. Isocontour lines are drawn in white. The model is visualized with electron density from the 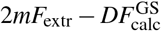 maps that produced the highest RSCCs for DW-Extrapolator and X8 in their respective grid scans. The electron density is contoured to 0.8*σ*. **(d)** Negative Log Likelihood landscape of ***θ*** = (*r, p*) given the experimental data. The maximum likelihood estimate 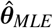 and default scan parameters ***θ***_Default Scan_ are plotted on the heatmap. The ligand binding-site is visualized with the difference map (purple/beige) and 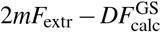 (gray) map for each of these recommended parameters. The maps are contoured as before.

Next, to evaluate the quality of these extrapolated structure factor amplitudes, we once again use Δ*R* and 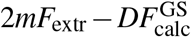 map RSCCs as quality metrics. DW-Extrapolator achieves Δ*R <* 0 for this dataset, showing that the extrapolation procedure results in structure factor amplitudes more consistent with the *holo* state than the *apo* state [Table III, Fig. 4(b)]. DW-Extrapolator’s *R*-factor values across our 2D parameter grid are around 10-15 percentage points lower than those produced in Xtrapol8’s scan of weighting schemes and occupancies [Fig. S4(b)]. DW-Extrapolator’s best 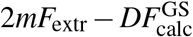 map, as measured by RSCC, displays strong electron density supporting the position and orientation of the drug fragment [Fig. 4(c)].

Finally, running DW-Extrapolator with both MLE and default scan parameters yields structure factor amplitudes that improve signal strength over the unextrapolated data, as measured by difference map and 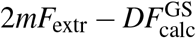 RSCCs, while achieving Δ*R <* 0 [Fig. 4(d)]. Xtrapol8’s recommended weighting and occupancy performs comparably by difference map RSCC, but produces 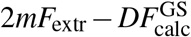 RSCC and *R*-factors that are around 10 percentage points lower and higher, respectively, than in the DW-Extrapolator cases [Tables II-IV, Fig. S4(d)].

### C. mEos4b Photoswitchable Fluorescent Protein

For our final test case, we use a chromophore excitation TR-X dataset in which crystals of mEos4b, a green-to-red photoswitchable fluorescent protein, were excited with 561-nm wavelength light to convert the protein from its *red-on* (i.e., red fluorescent) ground state to *red-off* (i.e., non-fluorescent) excited state^19,34^. These datasets were processed by the XDS^28^ software, which utilizes a Bayesian approach with a truncated normal posterior to estimate structure factor amplitudes from intensities, as proposed by French and Wilson^24^. Nevertheless, unlike Careless, XDS does not provide the parameters of its posterior distribution; instead, it provides only point estimates of *F*_**h**_, *σ* (*F*_**h**_). As such, we run DW-Extrapolator with the `–use_structure_factors` argument, which implements a method-of-moments procedure to estimate the parameters of a corresponding truncated normal posterior (see Supplementary Text IA).

Applying DW-Extrapolator to this dataset produces good performance across our metrics, attesting to the validity of this approach. The unextrapolated difference map shows little positive difference density corresponding to the chromophore’s excited *red-off* state [Fig. S5(a)]. The best difference map, as measured by RSCC, produced using DW-Extrapolator strengthens the excited-state density while also improving signal for other conformational changes near the chromophore [Fig. 5(a), Table II]. DW-Extrapolator also generates extrapolated structure factor amplitudes with lower *R*-factors than Xtrapol8 does for this dataset [Fig. S5(b)]. Still, as with the previous cases, the DW-Extrapolator structure factor amplitudes achieve Δ*R <* 0, confirming that extrapolation biases structure factor amplitudes towards being more consistent with the excited-state model [Fig. 5(b), Table III]. Further, DW-Extrapolator produces a best 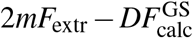 map that improves the unextrapolated map and features density for the chromophore’s excited conformation [Fig. 5(c), Table IV]. Finally, the MLE optimization and default scan procedures produce similar parameter estimates near (*r, p*) = (0.9, 0.3). Running DW-Extrapolator with these parameters results in improved map RSCCs over the unextrapolated data, while keeping *R*-factors around 0.25 [Fig. 5(d)]. Xtrapol8’s recommended weighting strategy and occupancy has lower map RSCCs, while yielding *R*-factors around 20 percentage points higher [Tables II-IV].

**FIG. 5.**
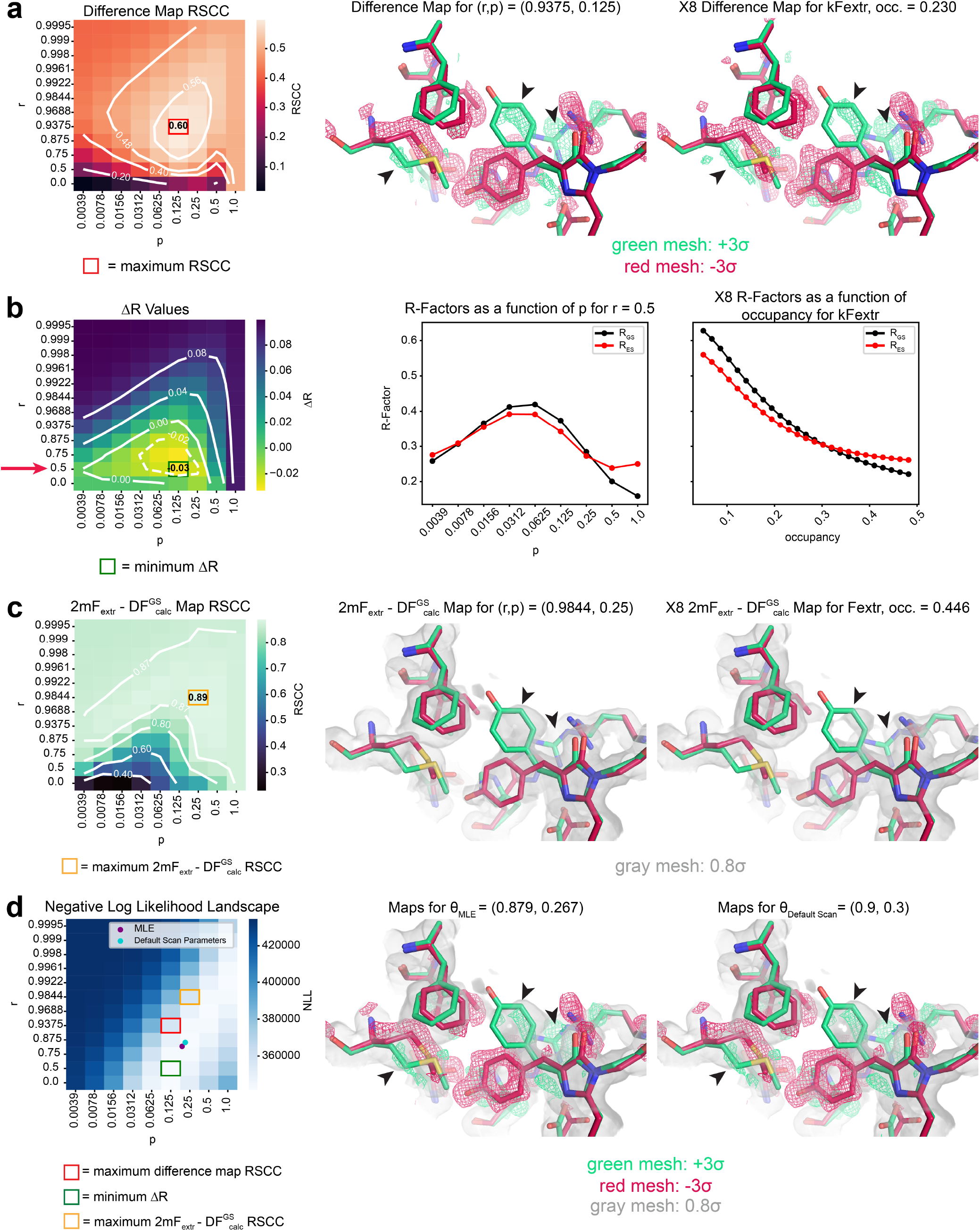
DW-Extrapolator results for the mEos4b chromophore excitation dataset. **(a)** Difference Map RSCC values within 10 Å of the mEos4b chromophore across a grid of (*r, p*). Isocontour lines are drawn in white. The mEos4b active site is visualized in *red-on* ground state (red) and *red-off* excited state (green). The difference density for the maps with the best RSCCs for the DW-Extrapolator and Xtrapol8 (X8) grid searches is contoured to ±3*σ*. Arrowheads indicate positive difference density corresponding to features of the excited state. **(b)** Δ*R*-factor values across the (*r, p*) grid, with isocontour lines drawn in white. The plot in the center fixes *r* = 0.5 and visualizes how *R*-factors change as a function of *p* for structure factor amplitudes produced by DW-Extrapolator. The plot on the right shows how *R*-factors change as a function of occupancy for *X*8’s kFextr extrapolation strategy. **(c)** 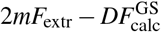 map RSCC values across (*r, p*) calculated within a 2 Å mask of the chromophore. Isocontour lines are drawn in white. The models are visualized with electron density from the 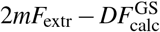 maps that produced the highest RSCCs for DW-Extrapolator and X8 across their grid scans. The electron density is contoured to 0.8*σ*. Arrowheads indicate map density corresponding to features of the excited state. **(d)** Negative Log Likelihood landscape of ***θ*** = (*r, p*) given the experimental data. The maximum likelihood estimate 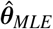 and default scan parameters ***θ***_Default Scan_ are plotted on the heatmap. The chromophore is visualized with the difference map (red/green) and 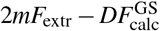 (gray) map for each of these recommended parameters. The maps are contoured as before. Arrowheads indicate positive difference density or 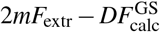 density corresponding to features of the excited state.

### D. Practical use

DW-Extrapolator is implemented as the `dw_extrapolate` command in the rs-booster package, and can handle datasets processed by a variety of crystallographic processing software. When run with 32 CPUs on a computing cluster, a single DW-Extrapolator run takes only a few minutes for each benchmark dataset [Table V]. To scan our 2D grid of (*r, p*) for our benchmarks, which contains 108 parameter combinations, we run parallel job arrays of single DW-Extrapolator runs, which take up to an hour, depending on computing resource constraints.

**TABLE V.** DW-Extrapolator Run Times.

| Dataset | Number of Reflections | Number of CPUs | Single Run | Default Scan | MLE Script |
| --- | --- | --- | --- | --- | --- |
| Photoactive Yellow Protein (PYP) | 12,138 | 32 | 2:08 | 22:08 | 3:38 |
| BAZ2B | 18,973 | 32 | 2:48 | 22:58 | 16:05 |
| mEos4b | 35,817 | 32 | 5:55 | 59:56 | 4:20 |

As a result, it may be more practical to run a 1D parameter sweep over *p*, as implemented in DW-Extrapolator’s default_scan mode. As executed by the command, this scan is not parallelized and takes up to 1 hour for the mEos4b dataset, which has 35,817 reflections. This time frame is similar to that required to run Xtrapol8 on a laptop computer, first in “fast-and-furious” mode to identify a range of plausible occupancies and then in “calm-and-curious” mode to test combinations of weighting schemes and occupancies [Table VI]. This process can be further expedited for DW-Extrapolator by running a 1D scan as a parallel job array and comparing the returned likelihood values. With tenfold less jobs than our 2D grid, running a 1D scan in parallel on a computing cluster takes around as long as a single job.

**TABLE VI.**
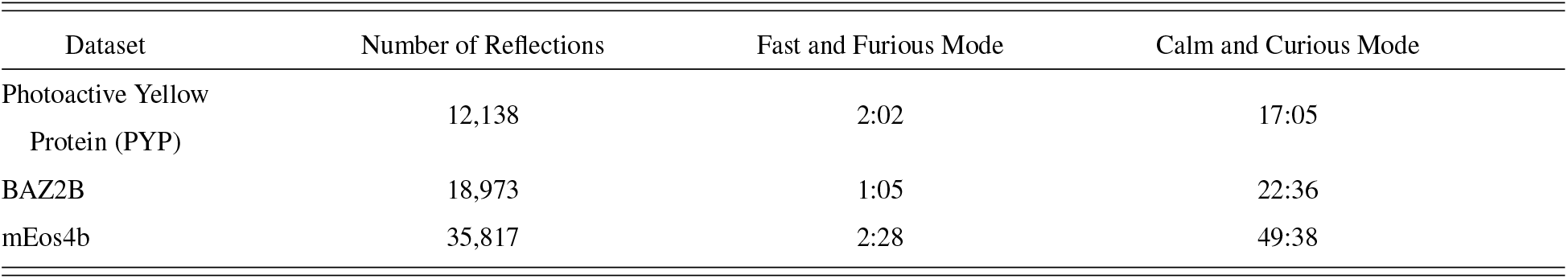
Xtrapol8 Run Times.

| Dataset | Number of Reflections | Fast and Furious Mode | Calm and Curious Mode |
| --- | --- | --- | --- |
| Photoactive Yellow Protein (PYP) | 12,138 | 2:02 | 17:05 |
| BAZ2B | 18,973 | 1:05 | 22:36 |
| mEos4b | 35,817 | 2:28 | 49:38 |

As illustrated in the above examples, the 1D scan reliably performs well. Although our 2D grid contains parameter combinations that outperform the default scan recommendation in each metric, we find that our three metrics are not perfectly correlated [Figs. 3-5(a)-(c)]. As such, in extrapolating structure factors for refinement, some tradeoff between signal amplification and noise suppression must be made. This tradeoff can be mediated by our statistical model’s likelihood function. Indeed, the default scan’s likelihood-based parameter recommendations are often comparable to the 2D grid’s optimum across multiple metrics.

For datasets in which the expected correlation *r* is unclear and for which the default scan doesn’t produce expected results, it may be helpful to run mle_dw_extrapolate to find the parameters 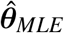 that achieve maximum likelihood in all of parameter space. Running this procedure would only add a few minutes to the data processing pipeline [Table V]. We note, however, that fixing *r* = 0.9 and then optimizing for *p* tends to produce slightly better results than a 2D optimization in our benchmark examples [Tables II-IV]. Given that the NLL landscape’s valley only enables optimization of the combination of parameters *r* and *p*, we recommend fixing *r* at a reasonable value as a first step.

## IV. DISCUSSION

Traditional structure factor extrapolation has succeeded in identifying features of the electron density corresponding to weakly populated excited states in perturbative crystallography experiments^1,4,6,14,41^. Using these extrapolated structure factor amplitudes for structure refinement, however, is challenging and produces models with poor *R*-factors due to the amplification of experimental errors and the assumption of equal phases between datasets^16,42^. Given that a major goal of perturbative crystallography experiments is to refine an excited-state structure, the shortcomings of traditional extrapolation are a significant constraint for the field.

The double-Wilson Extrapolator method begins to address these limitations. At its best, DW-Extrapolator performs appreciably better than traditional extrapolation: it better captures the details of the excited state, while the effects of error amplification are dampened by the double-Wilson prior’s correlation structure. The method’s Bayesian framework also allows for maximum-likelihood estimation of model parameters rather than the use of various, possibly conflicting, map and model metrics to select extrapolation parameters after the fact^4,19^. The use of a two-parameter model is, however, a clear simplification: the correlation between ground- and excited-state structure factors is almost certainly resolution-dependent^23^—we intend to address this in future work.

DW-Extrapolator is part of a growing body of work that adopts a Bayesian approach to infer a variety of estimands in the crystallographic data processing pipeline^21,29,43,44^. A natural question, then, is the extent to which these methods can be used in conjunction with each other. In our analysis of the PYP test case, we demonstrate that DW-Extrapolator can meaningfully amplify signal in datasets processed by Careless, which already uses variational Bayesian inference to estimate structure factor amplitudes^29^. A recent extension of Careless incorporates the double-Wilson prior into its statistical structure^21^. When we apply DW-Extrapolator to the same PYP dataset processed by Careless with the double-Wilson prior, however, we find no added benefit of using a double-Wilson prior twice [see Supplementary Text II]. As such, how statistical frameworks for different crystallographic data processing steps are best combined remains an open question.

## ACKNOWLEDGMENTS

We thank Drs. Alisia Fadini (Columbia University), Thomas J. Lane (DESY), Randy J. Read (University of Cambridge), Elke de Zitter and Jacques-Philippe Colletier (IBS, Grenoble), and Kevin Dalton (SLAC National Lab) for stimulating discussions. This work was supported by the Harvard College Research Program (to A.K.C.), National Science Foundation Graduate Research Fellowship grant DGE2140743 (to H.K.W.), and National Institutes of Health (NIH) grant DP2-GM141000 (to D.R.H.).

## DATA AVAILABILITY STATEMENT

The data and models used in this study are available from the Protein Data Bank under the following entries: 6GP0, 6GP1, and 1TS0. The PYP dataset was downloaded from Zenodo at 10.5281/zenodo.6408749. The BAZ2B dataset and models were downloaded from Zenodo at 10.5281/zenodo.48768. The extrapolated datasets and Jupyter notebooks used for the analyses and figures are available on Zenodo under accession code 10.5281/zenodo.19473677.

