## Supplementary Information for "Inferring structure factors of weakly populated excited states in perturbative crystallography experiments"

Andrew K. Choe,[1] Harrison K. Wang,[1,2] and Doeke R. Hekstra*[1,3]
[1)] *Department of Molecular and Cellular Biology, Harvard University, Cambridge, MA 02138*
[2)] *Graduate Program in Biophysics, Harvard University, Boston, MA 02115*
[3)] *John A. Paulson School of Engineering and Applied Sciences, Harvard University, Cambridge, MA 02138*

(*Electronic mail:)

(Dated: 16 April 2026)

**This file includes:**

1. Supplementary Text
2. Figures S1-S6
3. Supplementary References

### I. INFERRING EXCITED STATE AMPLITUDES

Double-Wilson (DW) Extrapolator can take as data either structure factor amplitudes or integrated intensities, depending on the crystallographic processing software used. The likelihood functions used in our Bayesian inference procedure differ across these two cases.

#### A. Using Structure Factor Amplitudes

Some merging and scaling software convert integrated intensities to structure factor amplitudes using a truncated normal posterior distribution, as introduced by French and Wilson[1]. As such, DW-Extrapolator uses a truncated normal likelihood function when working with data processed by these programs. Commonly used examples of software using this method include Careless[2] and XDS[3].

##### 1. Careless outputs

Conveniently, Careless[2] provides the parameters of the truncated normal posterior for the distribution of the true structure factor amplitude $F_{\mathbf{h}}$ for each Miller Index $\mathbf{h}$ given the corresponding observed diffraction data. For each observed reflection $\mathbf{h}$, Careless outputs the location ($\mu_{\mathbf{h}}$), scale ($\sigma_{\mathbf{h}}$), lower cut-point ($a = 10^{-32}$), and upper cut-point ($b = 10^{10}$) parameters for the posterior. Suppressing the cut-points in notation since they do not vary across reflections, let $f(F \mid \mu_{\mathbf{h}}, \sigma_{\mathbf{h}})$ be the truncated normal probability density function (PDF) associated with the posterior distribution of each reflection $\mathbf{h}$. The posterior likelihood inferred by DW-Extrapolator then becomes:

$$\begin{aligned}
P(\boldsymbol{E}_{\mathbf{h}}^{\mathrm{GS}}, \boldsymbol{E}_{\mathbf{h}}^{\mathrm{ES}} \mid E_{\mathbf{h}}^{\mathrm{off}}, E_{\mathbf{h}}^{\mathrm{on}}, \boldsymbol{\theta}) &\propto P(\boldsymbol{E}_{\mathbf{h}}^{\mathrm{GS}}, \boldsymbol{E}_{\mathbf{h}}^{\mathrm{ES}} \mid \boldsymbol{\theta}) \times P(E_{\mathbf{h}}^{\mathrm{off}}, E_{\mathbf{h}}^{\mathrm{on}} \mid \boldsymbol{E}_{\mathbf{h}}^{\mathrm{GS}}, \boldsymbol{E}_{\mathbf{h}}^{\mathrm{ES}}, \boldsymbol{\theta}) \\
&\approx P(\boldsymbol{E}_{\mathbf{h}}^{\mathrm{GS}}, \boldsymbol{E}_{\mathbf{h}}^{\mathrm{ES}} \mid \boldsymbol{\theta}) \times P(E_{\mathbf{h}}^{\mathrm{off}} \mid \boldsymbol{E}_{\mathbf{h}}^{\mathrm{GS}}, \boldsymbol{\theta}) \times P(E_{\mathbf{h}}^{\mathrm{on}} \mid \boldsymbol{E}_{\mathbf{h}}^{\mathrm{ON}}, \boldsymbol{\theta}) \\
&= P(\boldsymbol{E}_{\mathbf{h}}^{\mathrm{GS}}, \boldsymbol{E}_{\mathbf{h}}^{\mathrm{ES}} \mid \boldsymbol{\theta}) \times \frac{P(\boldsymbol{E}_{\mathbf{h}}^{\mathrm{GS}} \mid E_{\mathbf{h}}^{\mathrm{off}}, \boldsymbol{\theta}) \times P(E_{\mathbf{h}}^{\mathrm{off}} \mid \boldsymbol{\theta})}{P(\boldsymbol{E}_{\mathbf{h}}^{\mathrm{GS}} \mid \boldsymbol{\theta})} \\
&\qquad \times \frac{P(\boldsymbol{E}_{\mathbf{h}}^{\mathrm{ON}} \mid E_{\mathbf{h}}^{\mathrm{on}}, \boldsymbol{\theta}) \times P(E_{\mathbf{h}}^{\mathrm{on}} \mid \boldsymbol{\theta})}{P(\boldsymbol{E}_{\mathbf{h}}^{\mathrm{ON}} \mid \boldsymbol{\theta})} \\
&\approx P(\boldsymbol{E}_{\mathbf{h}}^{\mathrm{GS}}, \boldsymbol{E}_{\mathbf{h}}^{\mathrm{ON}} \mid \boldsymbol{\theta}) \times f^{\mathrm{off}}\left(|\boldsymbol{E}_{\mathbf{h}}^{\mathrm{GS}}| \times \sqrt{\varepsilon_{\mathbf{h}} \Sigma_{\mathbf{h}}} \mid \mu_{\mathbf{h}}^{\mathrm{off}}, \sigma_{\mathbf{h}}^{\mathrm{off}}\right) \\
&\qquad \times f^{\mathrm{on}}\left(|\boldsymbol{E}_{\mathbf{h}}^{\mathrm{ON}}| \times \sqrt{\varepsilon_{\mathbf{h}} \Sigma_{\mathbf{h}}} \mid \mu_{\mathbf{h}}^{\mathrm{on}}, \sigma_{\mathbf{h}}^{\mathrm{on}}\right), \qquad (1)
\end{aligned}$$

where $\boldsymbol{E}_{\mathbf{h}}^{\mathrm{ON}} = (1-p)\boldsymbol{E}_{\mathbf{h}}^{\mathrm{GS}} + p\boldsymbol{E}_{\mathbf{h}}^{\mathrm{ES}}$. Further, we go from the penultimate to final line above by assuming that the prior probabilities of the observed amplitudes ($E_{\mathbf{h}}^{\mathrm{off}}$ and $E_{\mathbf{h}}^{\mathrm{on}}$) are roughly equal to the prior probabilities of their true complex values under the double-Wilson prior ($\boldsymbol{E}_{\mathbf{h}}^{\mathrm{GS}}$ and $\boldsymbol{E}_{\mathbf{h}}^{\mathrm{ON}}$, respectively).

##### 2. Non-Careless outputs (use_structure_factors mode)

Other merging and scaling programs that utilize a truncated normal posterior do not explicitly give distribution parameters as Careless does. Instead, they usually only output point estimates for $(F_{\mathbf{h}}, \sigma(F_{\mathbf{h}}))$. Using a method of moments approach, we can treat these point estimates as the mean and standard deviation, respectively, of the underlying truncated normal distribution.

Suppressing the Miller index, let $F \sim \mathrm{TruncatedNormal}(\mu, \sigma, a, b)$. Define the shape parameters of $F$ as[4]:

$$\alpha = \frac{a-\mu}{\sigma} \qquad (2)$$

$$\beta = \frac{b-\mu}{\sigma}, \qquad (3)$$

where we set $a = 10^{-32}, b = 10^{10}$ as in the Careless case. Let $\Phi$ denote the standard normal cumulative distribution function (CDF) and $\varphi$ denote the standard normal PDF. The expectation of $F$ is[4]:

$$\mathbb{E}[F] = \mu - \sigma \times \frac{\varphi(\beta) - \varphi(\alpha)}{\Phi(\beta) - \Phi(\alpha)} \tag{4}$$

The variance of $F$ is[4]:

$$\mathrm{Var}(F) = \sigma^2 \times \left[ 1 - \frac{\beta\varphi(\beta) - \alpha\varphi(\alpha)}{\Phi(\beta) - \Phi(\alpha)} - \left( \frac{\varphi(\beta) - \varphi(\alpha)}{\Phi(\beta) - \Phi(\alpha)} \right)^2 \right] \tag{5}$$

For fixed cut-points $(a, b)$, note that $\mu$ and $\sigma$ can be expressed as functions of only the shape parameters:

$$\sigma = \frac{b-a}{\beta - \alpha} = f(\alpha, \beta) \tag{6}$$

$$\mu = a - \sigma\alpha = b - \sigma\beta = g(\alpha, \beta) \tag{7}$$

Let $\bar{F}, \bar{\sigma}(F)$ be the observed structure factor amplitude and its associated error, respectively. Treat these observations as the sample moments of $F$. Then, adopting a method of moments approach, we set the sample moments equal to their true distribution values:

$$\begin{cases} \bar{F} & = \mathbb{E}[F] = g(\alpha, \beta) \\ \bar{\sigma}(F) & = \mathrm{Var}(F) = f(\alpha, \beta) \end{cases} \tag{8}$$

The above in Eq. (8) is a system of two equations in two unknowns $(\alpha, \beta)$. No closed form solution exists due to the normal CDF ($\Phi$), so we numerically estimate a solution $(\hat{\alpha}, \hat{\beta})$. We can transform these estimates using Eqs. (6), (7) to get estimates of $(\hat{\mu}, \hat{\sigma})$.

Running this procedure for all Miller indices $\mathbf{h}$ then yields truncated normal parameter estimates for each reflection. With these parameters in hand, we can then calculate the posterior likelihood in the same way as described in Eq. (1).

### B. Using Intensities (use_intensities mode)

Other merging and scaling programs use physically, not statistically, motivated methods for estimating structure factor amplitudes. In such cases, DW-Extrapolator works directly with integrated intensities. Following the statistical model introduced by French and Wilson[1], we assume that observed intensities $I_{\mathbf{h}}$ are Normally distributed about their true, necessarily positive mean values $J_{\mathbf{h}}$:

$$I_{\mathbf{h}} \sim \mathcal{N}(J_{\mathbf{h}}, \sigma_{\mathbf{h}}^2) \tag{9}$$

We will assume the experimentally estimated errors for the intensities $\sigma(I_{\mathbf{h}})$ to be good approximations for the $\sigma_{\mathbf{h}}$ term in (9). Further, note that diffraction intensities are proportional to square of the associated structure factor amplitude[5]:

$$J_{\mathbf{h}} = \varepsilon_{\mathbf{h}} \Sigma_{\mathbf{h}} |\boldsymbol{E}_{\mathbf{h}}|^2 \tag{10}$$

Given this statistical model on intensities, the posterior inferred by DW-Extrapolator then becomes:

$$\begin{aligned} P(\boldsymbol{E}_{\mathbf{h}}^{\mathrm{GS}}, \boldsymbol{E}_{\mathbf{h}}^{\mathrm{ES}} \mid I_{\mathbf{h}}^{\mathrm{off}}, I_{\mathbf{h}}^{\mathrm{on}}, \boldsymbol{\theta}) &\propto P(\boldsymbol{E}_{\mathbf{h}}^{\mathrm{GS}}, \boldsymbol{E}_{\mathbf{h}}^{\mathrm{ES}} \mid \boldsymbol{\theta}) \times P(I_{\mathbf{h}}^{\mathrm{off}}, I_{\mathbf{h}}^{\mathrm{on}} \mid \boldsymbol{E}_{\mathbf{h}}^{\mathrm{GS}}, \boldsymbol{E}_{\mathbf{h}}^{\mathrm{ES}}, \boldsymbol{\theta}) \\ &\approx P(\boldsymbol{E}_{\mathbf{h}}^{\mathrm{GS}}, \boldsymbol{E}_{\mathbf{h}}^{\mathrm{ES}} \mid \boldsymbol{\theta}) \times P(I_{\mathbf{h}}^{\mathrm{off}} \mid \boldsymbol{E}_{\mathbf{h}}^{\mathrm{GS}}, \boldsymbol{\theta}) \times P(I_{\mathbf{h}}^{\mathrm{on}} \mid \boldsymbol{E}_{\mathbf{h}}^{\mathrm{ON}}, \boldsymbol{\theta}) \\ &\propto P(\boldsymbol{E}_{\mathbf{h}}^{\mathrm{GS}}, \boldsymbol{E}_{\mathbf{h}}^{\mathrm{ES}} \mid \boldsymbol{\theta}) \times \varphi\left( \frac{I_{\mathbf{h}}^{\mathrm{off}} - \varepsilon_{\mathbf{h}} \Sigma_{\mathbf{h}} |\boldsymbol{E}_{\mathbf{h}}^{\mathrm{GS}}|^2}{\sigma(I_{\mathbf{h}}^{\mathrm{off}})} \right) \times \varphi\left( \frac{I_{\mathbf{h}}^{\mathrm{on}} - \varepsilon_{\mathbf{h}} \Sigma_{\mathbf{h}} |\boldsymbol{E}_{\mathbf{h}}^{\mathrm{ON}}|^2}{\sigma(I_{\mathbf{h}}^{\mathrm{on}})} \right), \end{aligned} \tag{11}$$

where $\boldsymbol{E}_{\mathbf{h}}^{\mathrm{ON}} = (1-p)\boldsymbol{E}_{\mathbf{h}}^{\mathrm{GS}} + p\boldsymbol{E}_{\mathbf{h}}^{\mathrm{ES}}$ as before and $\varphi$ is the standard normal PDF.

### II. DW-EXTRAPOLATOR OF DATA PROCESSED BY CARELESS WITH THE DOUBLE-WILSON PRIOR

Recently, Hekstra *et al.* introduced the double-Wilson prior into the inference framework of the scaling software Careless[6]. Interested in how DW-Extrapolator would perform for data that had already seen the double-Wilson prior during scaling, we ran `dw_extrapolate` on the same PYP dataset, but now processed using Careless with the DW prior.

We observed that DW-Extrapolator did not significantly improve signal quality beyond Careless with the double-Wilson prior. Difference map and $2mF_{\text{extr}} - DF_{\text{calc}}^{\text{GS}}$ map real space correlation coefficients (RSCCs) were comparable between the extrapolated and unextrapolated versions [Tables S1, S2]. If anything, scaling with the double-Wilson prior appears to have improved the performance of traditional extrapolation with Xtrapol8 while slightly diminishing the performance of DW-Extrapolator. A minor improvement was observed in the form of lower $R$-factors [Table S3].

TABLE S1. Maximum PYP Dataset Difference Map RSCC Values across Careless Versions

| Processing Software | Unextrapolated RSCC | DW-Extrapolator 2D Grid Max. | Xtrapol8 Grid Max. |
|---|---|---|---|
| Careless | 0.38 | 0.45 | 0.42 |
| Careless with DW prior | 0.49 | 0.48 | 0.49 |

TABLE S2. Maximum PYP Dataset $2mF_{\text{extr}} - DF_{\text{calc}}^{\text{GS}}$ Map RSCC Values across Careless Versions

| Processing Software | Unextrapolated RSCC | DW-Extrapolator 2D Grid Max. | Xtrapol8 Grid Max. |
|---|---|---|---|
| Careless | 0.694 | 0.734 | 0.656 |
| Careless with DW prior | 0.683 | 0.711 | 0.764 |

TABLE S3. $R_{\text{GS}}$ and $R_{\text{ES}}$ values at the minimum $\Delta R$ for PYP Dataset

| Processing Software | DW-Extrapolator $R_{\text{GS}}$ | DW-Extrapolator $R_{\text{ES}}$ | Xtrapol8 $R_{\text{GS}}$ | Xtrapol8 $R_{\text{ES}}$ |
|---|---|---|---|---|
| Careless | 0.40 | 0.38 | 0.57 | 0.53 |
| Careless with DW prior | 0.37 | 0.35 | 0.65 | 0.61 |

### SUPPLEMENTARY FIGURES

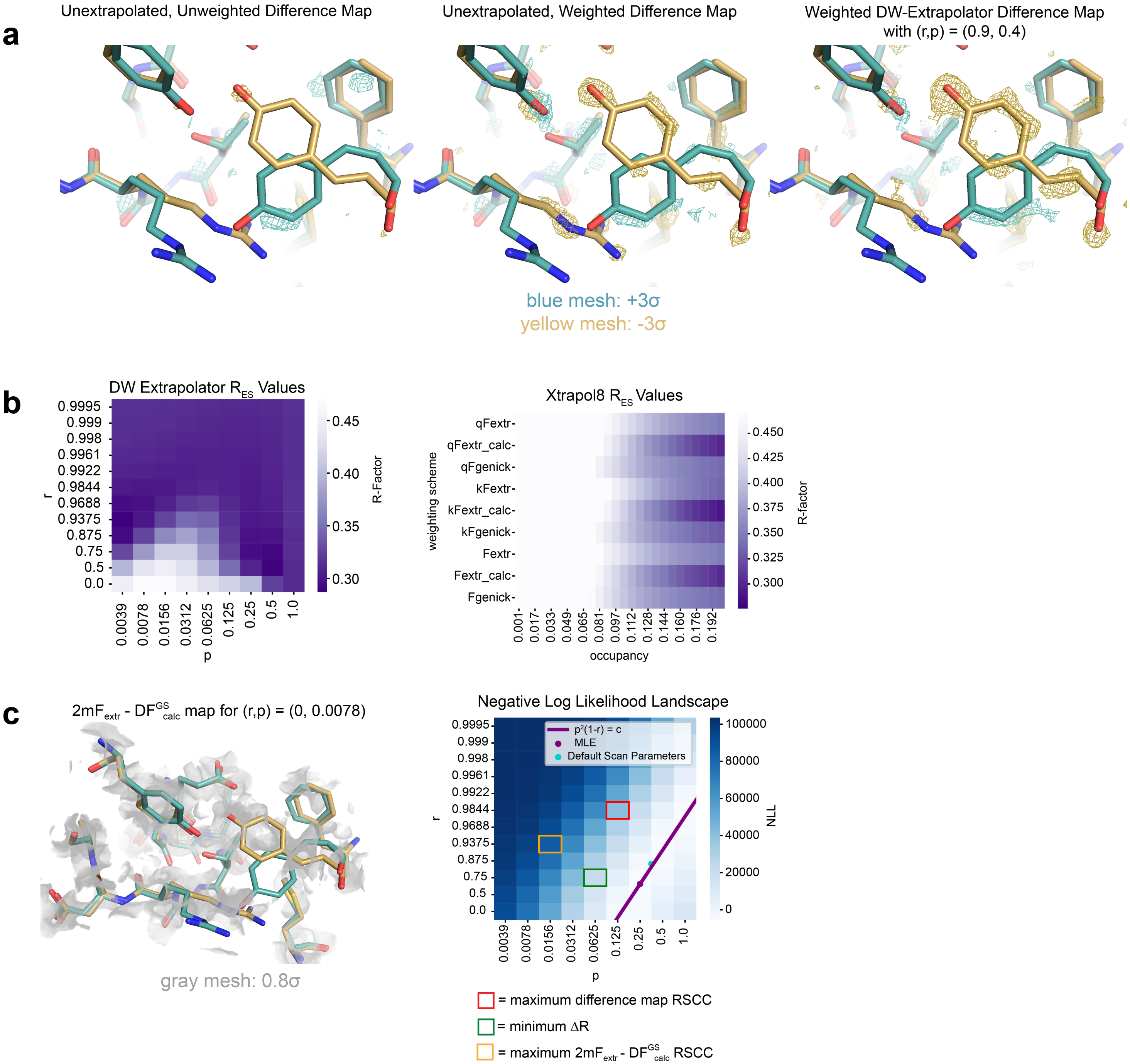


FIG. S1. *Caption on next page.*

FIG. S1. **Assessing DW-Extrapolator's performance on the PYP dataset.** **(a)** Comparison of unweighted and weighted $F^{\text{on}} - F^{\text{off}}$ difference maps for the unextrapolated structure factors. The ground state model is shown in yellow, and the excited state model is shown in teal. Weighting strengthens the difference density signal of the PYP chromophore isomerization. Still, even with the weighting scheme, the difference density is weak and discontinuous, making it difficult to see details of the conformational change. A weighted difference map produced with extrapolated structure factors features much stronger difference density for the chromophore's ground and excited states. **(b)** Comparison between excited state $R$-factors for DW-Extrapolator and Xtrapol8. The color scales for the two heatmaps are the same. Overall, DW-Extrapolator produces structure factors with lower $R$-factors, suggesting that it is more robust to phase and measurement errors. **(c)** $2mF_{\text{extr}} - DF_{\text{calc}}^{\text{GS}}$ map produced by DW-Extrapolator with the worst RSCC. This visualization confirms that low $2mF_{\text{extr}} - DF_{\text{calc}}^{\text{GS}}$ map RSCC values correspond to structure factors that poorly fit the model. **(d)** Negative Log Likelihood (NLL) landscape for $(r, p)$ given the PYP dataset. The valley where the NLL achieves its minimum values is closely approximated by the curve $p^2(1-r) = c$, where $c = \hat{p}_{MLE}^2(1-\hat{r}_{MLE})$.

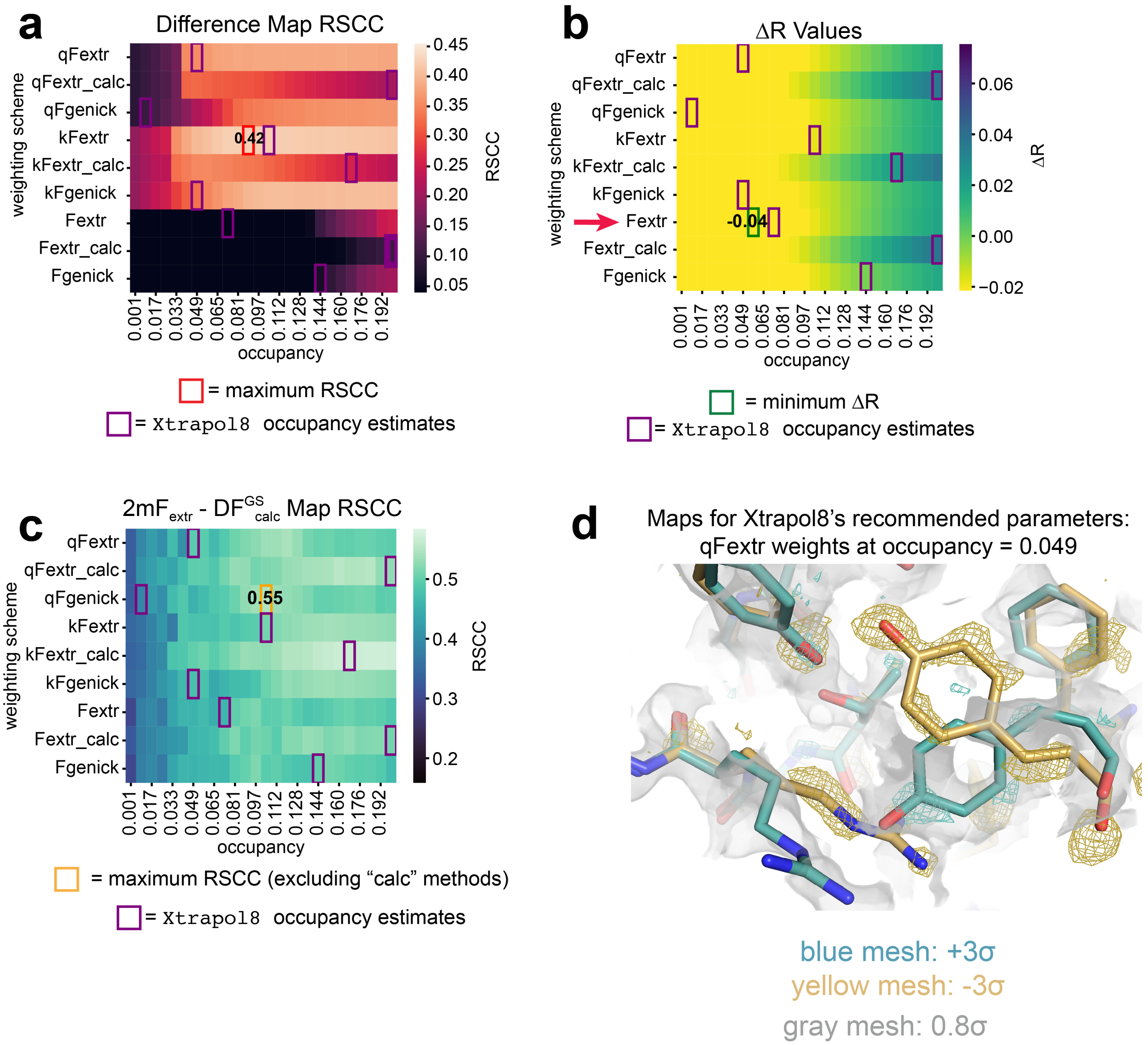


FIG. S2. **Xtrapol8 PYP benchmark.** For ease of comparison, all colormaps are on the same scale as the corresponding plot in Fig. 3 of the main text. **(a)** Difference Map RSCC values within 10 Å of the chromophore across a grid of occupancies and Xtrapol8's extrapolation strategies. **(b)** $\Delta R$-factor values across the extrapolation strategies. **(c)** $2mF_{\mathrm{extr}} - DF^{\mathrm{GS}}_{\mathrm{calc}}$ map RSCC values computed in a 2 Å mask around the chromophore. In consideration of the maximum, we exclude the $F_{\mathrm{calc}}$ methods, since these extrapolate from structure factors calculated from the ground state model, producing artificially low $R$-factors. The electron density is contoured to $0.8\sigma$. **(d)** Difference and $2mF_{\mathrm{extr}} - DF^{\mathrm{GS}}_{\mathrm{calc}}$ maps for the structure factors extrapolated using Xtrapol8's recommended weighting strategy and occupancy. The molecular model shows the PYP chromophore with the difference (teal/yellow) and $2mF_{\mathrm{extr}} - DF^{\mathrm{GS}}_{\mathrm{calc}}$ (gray) maps. The maps are contoured as in the main text.

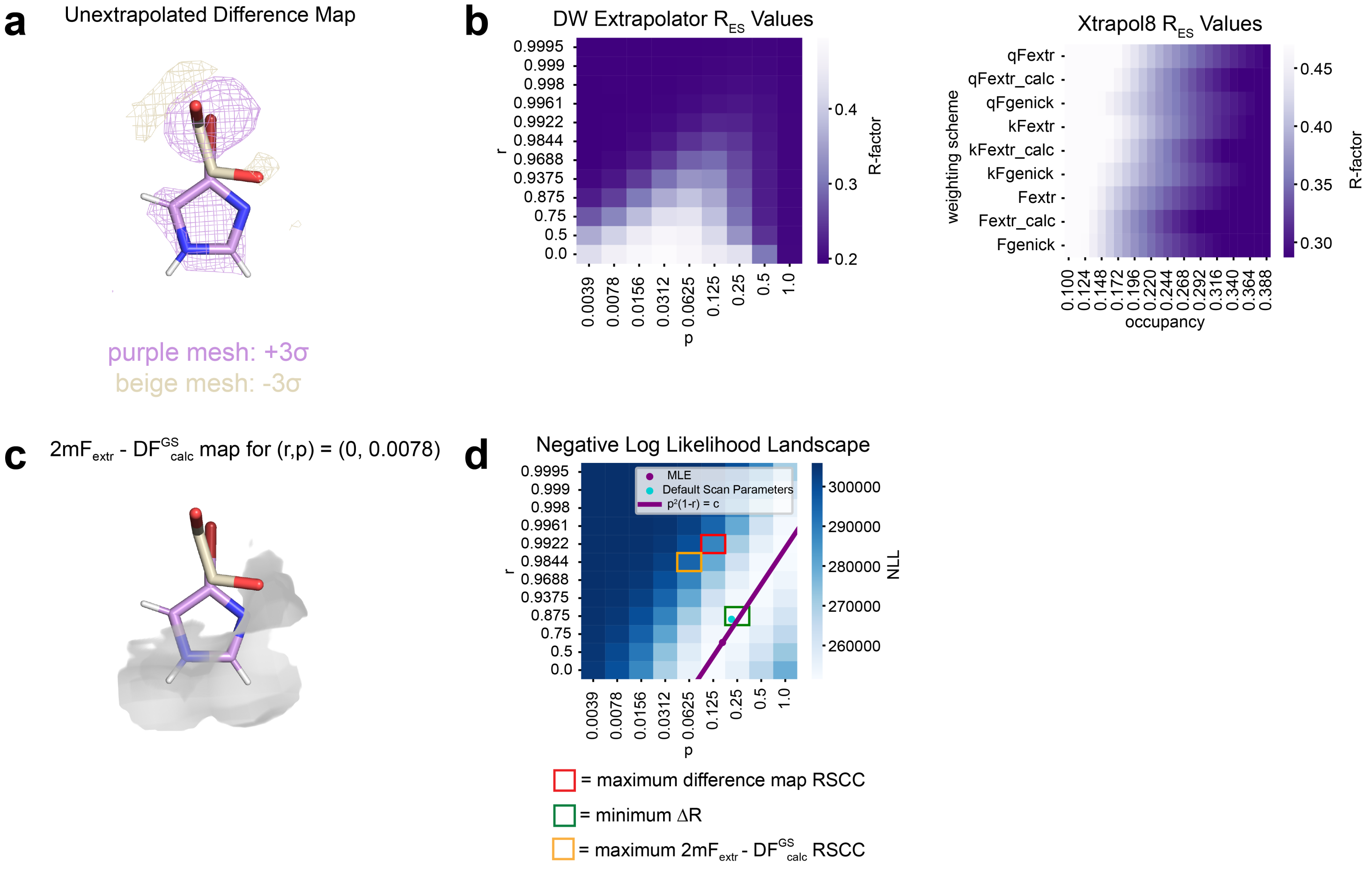


FIG. S3. **Assessing DW-Extrapolator's performance on the BAZ2B benchmark. (a)** Unextrapolated $F^{\text{on}} - F^{\text{off}}$ difference map with *apo* (beige) and *holo* (purple) state models. For this fragment screening dataset, the unextrapolated difference map displays appreciable difference density, yet all the features of the electron density are less pronounced than in the best DW-Extrapolator difference maps. **(b)** Comparison between $R_{\text{ES}}$ $R$-factor values for DW Extrapolation and Xtrapol8. DW-Extrapolator produces structure factors with lower $R$-factors, indicating that they are more accurate. **(c)** $2mF_{\text{extr}} - DF^{\text{GS}}_{\text{calc}}$ map produced by DW-Extrapolator with the worst RSCC. This visualization confirms that low $2mF_{\text{extr}} - DF^{\text{GS}}_{\text{calc}}$ map RSCC values correspond to structure factors that poorly fit the model. **(d)** Negative Log Likelihood (NLL) landscape for $(r, p)$ given the BAZ2B dataset. The valley where the NLL achieves its minimum values is closely approximated by the curve $p^2(1-r) = c$, where $c = \hat{p}^2_{MLE}(1-\hat{r}_{MLE})$.

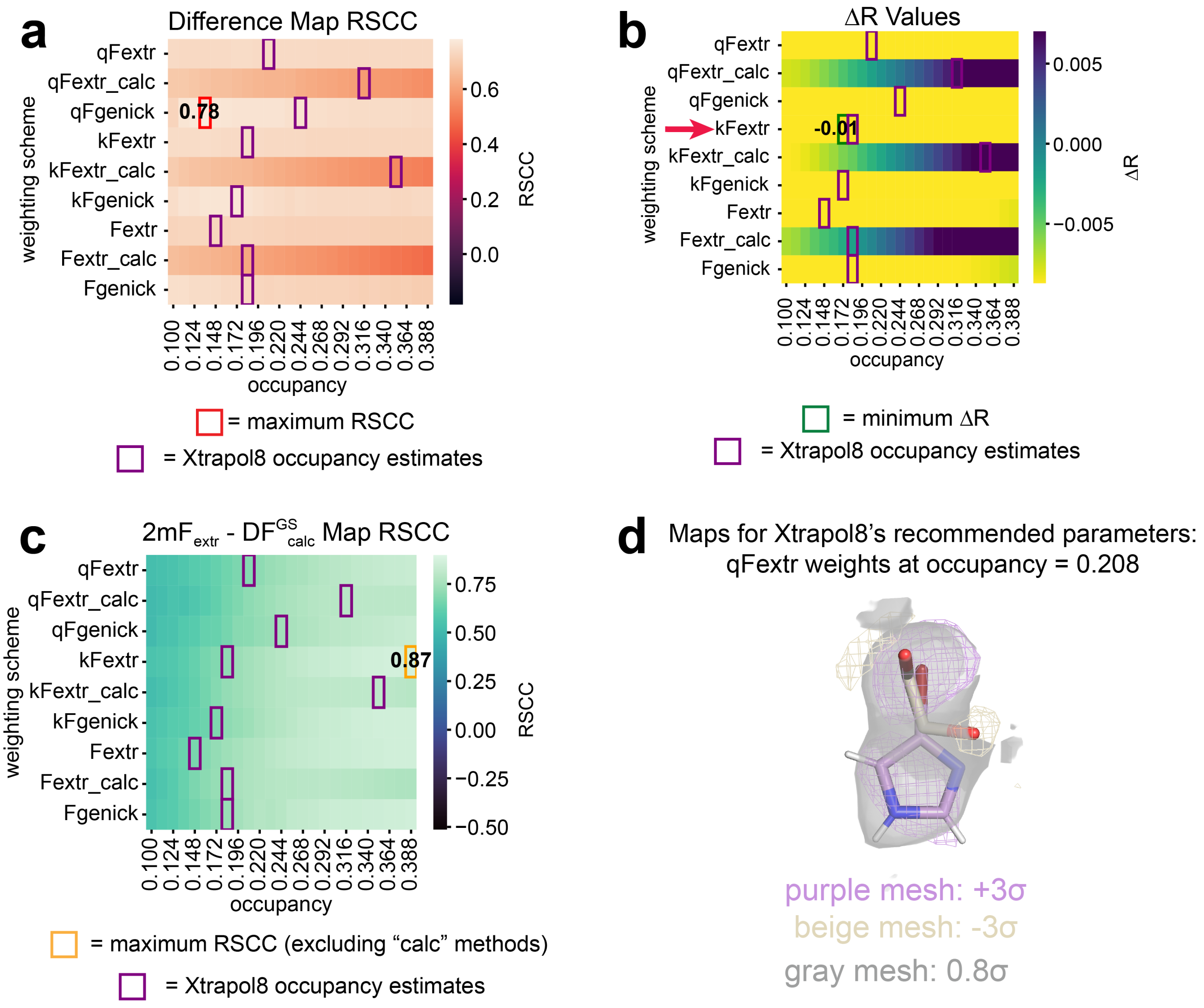


FIG. S4. **Xtrapol8 BAZ2B fragment screen benchmark.** All colormaps are on the same scale as the corresponding plot in Fig. 4 of the main text. **(a)** Difference Map RSCC values for a 2 Å mask around the ligand molecule. **(b)** $\Delta R$-factor values across the extrapolation strategies. **(c)** $2mF_{\text{extr}} - DF_{\text{calc}}^{\text{GS}}$ map RSCC values calculated for a 2 Å mask around the ligand. As before, we exclude the $F_{\text{calc}}$ methods from the maximum calculation, since these approaches extrapolate from structure factors calculated from the ground state model. The electron density is contoured to $0.8\sigma$. **(d)** Difference (purple/beige) and $2mF_{\text{extr}} - DF_{\text{calc}}^{\text{GS}}$ (gray) maps for the structure factors extrapolated using Xtrapol8's recommended weighting strategy and occupancy. The maps are contoured as before.

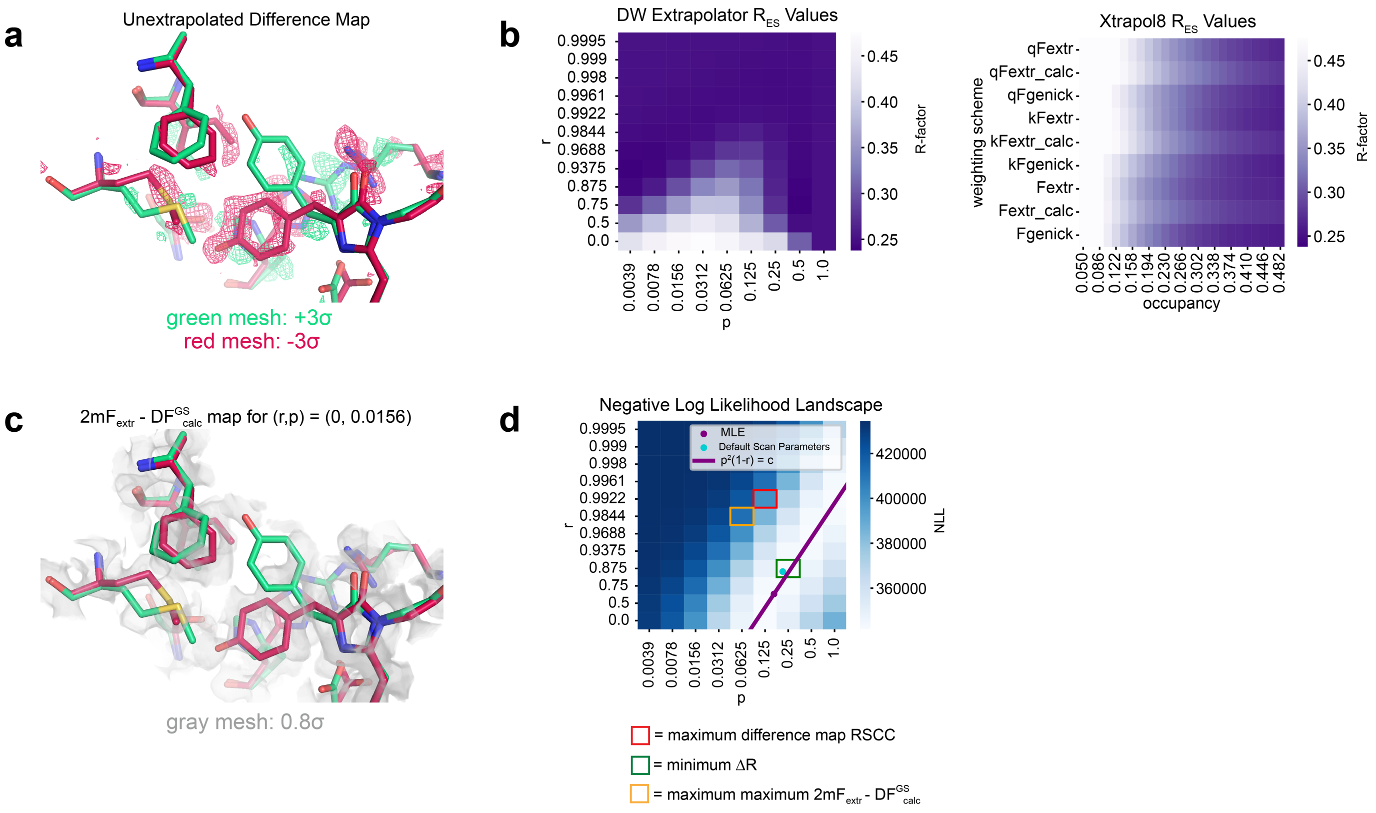


FIG. S5. Assessing DW-Extrapolator's performance on the mEos4b benchmark. **(a)** Unextrapolated $F^{\text{on}} - F^{\text{off}}$ difference map with *red-on* (red) and *red-off* (green) state models. The unextrapolated density for the rotamer flips near the chromophore are weakly supported by the difference density. **(b)** Comparison between $R_{\text{ES}}$ $R$-factor values for DW-Extrapolator and Xtrapol8. DW-Extrapolator produces structure factors with lower $R$-factors, suggesting that it is more robust to phase and measurement errors. **(c)** $2mF_{\text{extr}} - DF^{\text{GS}}_{\text{calc}}$ map produced by DW-Extrapolator with the worst RSCC. This visualization confirms that low $2mF_{\text{extr}} - DF^{\text{GS}}_{\text{calc}}$ map RSCC values correspond to structure factors that poorly fit the model. **(d)** Negative Log Likelihood (NLL) landscape for $(r, p)$ given the mEos4b dataset. The valley where the NLL achieves its minimum values is closely approximated by the curve $p^2(1-r) = c$, where $c = \hat{p}^2_{MLE}(1 - \hat{r}_{MLE})$.

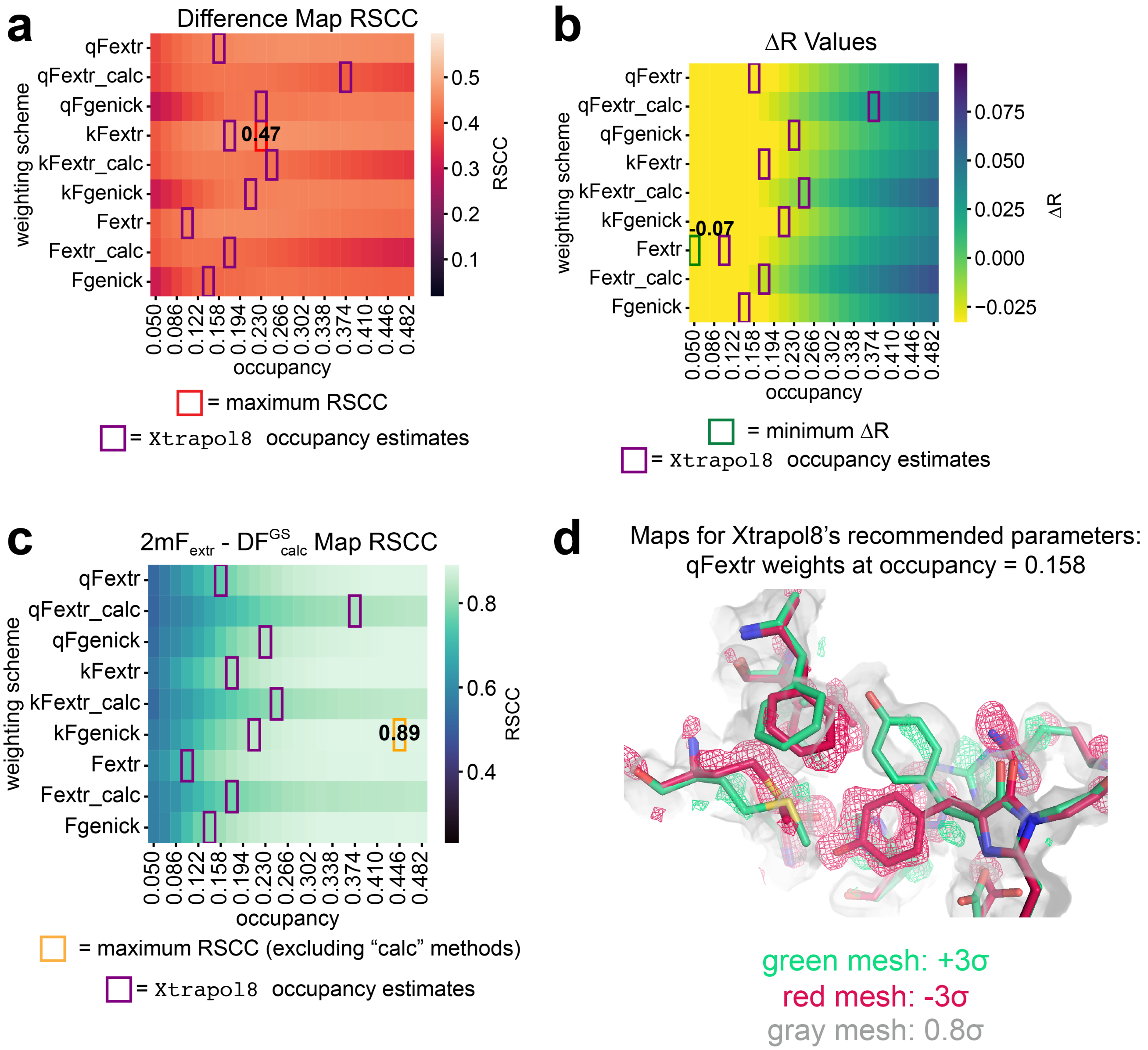


FIG. S6. Xtrapol8 mEos4b benchmark. All colormaps are on the same scale as the corresponding plot in Fig. 5 of the main text. **(a)** Difference Map RSCC values within 10 Å of the chromophore. **(b)** $\Delta R$-factor values across the extrapolation strategies. **(c)** $2mF_{\mathrm{extr}} - DF^{\mathrm{GS}}_{\mathrm{calc}}$ map RSCC values across extrapolation strategies. As before, we exclude the $F_{\mathrm{calc}}$ methods from the maximum calculation, since these approaches extrapolate from structure factors calculated from the ground state model. The electron density is contoured to $0.8\sigma$. **(d)** Difference (green) and $2mF_{\mathrm{extr}} - DF^{\mathrm{GS}}_{\mathrm{calc}}$ (gray) maps for the structure factors extrapolated using Xtrapol8's recommended weighting strategy and occupancy. The maps are contoured as before.
